# SlKIX8 and SlKIX9 are negative regulators of leaf and fruit growth in tomato

**DOI:** 10.1101/2020.02.07.938977

**Authors:** Gwen Swinnen, Jean-Philippe Mauxion, Alexandra Baekelandt, Rebecca De Clercq, Jan Van Doorsselaere, Dirk Inzé, Nathalie Gonzalez, Alain Goossens, Laurens Pauwels

**Author notes:** current address: University of Lausanne, Center for Integrative Genomics, 1015 Lausanne, Switzerland (Gwen Swinnen). equally contributed (Gwen Swinnen, Jean-Philippe Mauxion). senior authors (Alain Goossens, Laurens Pauwels). corresponding author: Laurens Pauwels, VIB-UGent Center for Plant Systems Biology, Technologiepark 71, B-9052 Gent (Belgium). **Author Contribution:** G.S., J-P.M., A.B., L.P., N.G., and A.G. designed the experiments. G.S., J-P.M., A.B., R.D.C., and J.V.D. performed the experiments. G.S., J-P.M., A.B., L.P., N.G., and A.G. analyzed the data. G.S. wrote the article and J-P.M., A.B., L.P., N.G., and A.G. contributed to the writing and revised the manuscript. N.G., D.I., A.G. and L.P. supervised the project. L.P. agrees to serve as the author responsible for contact and ensures communication.

## Abstract

Plant organ size and shape are major agronomic traits that depend on cell division and expansion, which are both regulated by complex gene networks. In several eudicot species belonging to the rosid clade, organ growth is controlled by a repressor complex consisting of PEAPOD (PPD) and KINASE-INDUCIBLE DOMAIN INTERACTING (KIX) proteins. The role of these proteins in asterids, which together with the rosids constitute most of the core eudicot species, is still unknown. We used CRISPR-Cas9 genome editing to target *SlKIX8* and *SlKIX9* in the asterid model species tomato (*Solanum lycopersicum*) and analyzed loss-of-function phenotypes. We found that loss of function of *SlKIX8* and *SlKIX9* led to the production of enlarged, dome-shaped leaves and that these leaves exhibited increased expression of putative SlPPD target genes. Unexpectedly, *kix8 kix9* mutants carried enlarged fruits with increased pericarp thickness due to cell expansion. At the molecular level, protein interaction assays indicated that SlKIX8 and SlKIX9 act as adaptors between the SlPPD and SlTOPLESS co-repressor proteins. Our results show that KIX8 and KIX9 are regulators of organ growth in asterids and can provide strategies to improve important traits in produce such as thickness of the fruit flesh.

**One sentence summary:** Two transcriptional repressors negatively regulate organ growth in tomato with loss-of-function lines producing enlarged fruits due to an appearance of more expanded cells in the fruit flesh.

## Introduction

Plants come in all shapes and sizes, yet these agronomically important traits are remarkably uniform within a given species or variety. Not surprisingly, cell division and cell expansion, the underlying processes of organ development, are under tight genetic control (Gonzalez *et al.*, 2012; Hepworth & Lenhard, 2014; Kalve *et al.*, 2014; Vercruysse *et al.*, 2020). The different phases of leaf development, for instance, are regulated by complex gene networks (Gonzalez *et al.*, 2012; Hepworth & Lenhard, 2014; Vercruysse *et al.*, 2020). Leaf development consists of the emergence of a leaf primordium from the shoot apical meristem, followed by a period of primary cell division that transitions into a cell expansion phase, and a simultaneous phase of meristemoid division. In Arabidopsis (*Arabidopsis thaliana*) leaves, the shape of the primary cell cycle arrest front, which moves from tip to base as cells cease to divide, was reported to be regulated by the transcriptional regulator PEAPOD2 (AtPPD2) (Baekelandt *et al.*, 2018). Through its interaction with the adaptor protein NOVEL INTERACTOR OF JAZ (AtNINJA) (Supplemental Figure 1A), AtPPD2 can recruit the transcriptional co-repressor TOPLESS (AtTPL) and, thereby, control leaf flatness (Baekelandt *et al.*, 2018). In addition, AtPPD2 forms a transcriptional repressor complex with KINASE-INDUCIBLE DOMAIN INTERACTING 8 (AtKIX8)/AtKIX9 (Supplemental Figure 1A) (Gonzalez *et al.*, 2015), which can also recruit AtTPL, to limit the number of self-renewing asymmetric divisions that stem-cell like meristemoids can undergo before differentiating into stomatal guard cells (White, 2006; Gonzalez *et al.*, 2015). This way, the AtPPD2-AtKIX8/AtKIX9 repressor complex restricts leaf growth, thereby controlling both leaf shape and size (White, 2006; Gonzalez *et al.*, 2015). The functionalities of AtKIX8 and AtKIX9 are, thus, required for the repressive activity of AtPPD2 (Gonzalez *et al.*, 2015). Consequently, double *kix8-kix9* Arabidopsis knockout plants display increased transcript levels of AtPPD2 target genes and enlarged, dome-shaped leaves because of a prolonged period of meristemoid division, similar to Arabidopsis *ami-ppd* plants (Gonzalez *et al.*, 2015).

In Arabidopsis, the AtKIX8 and AtKIX9 proteins are known to interact with both AtPPD1 and AtPPD2 through their distinguishing N-terminal PPD domain (Supplemental Figure 1A) (Bai *et al.*, 2011; Gonzalez *et al.*, 2015). Together with AtTIFY8 and the JASMONATE ZIM DOMAIN (AtJAZ) proteins, AtPPD1 and AtPPD2 belong to class II of the TIFY protein family (Vanholme *et al.*, 2007; Bai *et al.*, 2011) and are characterized by the presence of a conserved TIF[F/Y]XG motif. This motif resides within the ZINC-FINGER PROTEIN EXPRESSED IN INFLORESCENCE MERISTEM (ZIM) domain that mediates the interaction of class II TIFY proteins with AtNINJA (Supplemental Figure 1A) (Chini *et al.*, 2009; Chung & Howe, 2009; Pauwels *et al.*, 2010; Baekelandt *et al.*, 2018). All class II proteins, except AtTIFY8, contain a C-terminal Jas domain (Vanholme *et al.*, 2007; Bai *et al.*, 2011). The Jas domain found in AtPPD proteins, however, is divergent from the Jas consensus motif in AtJAZ proteins (Bai *et al.*, 2011) that mediates the interaction of AtJAZ proteins with transcription factors such as AtMYC2 and the F-box protein CORONATINE INSENSITIVE 1 (AtCOI1) (Chini *et al.*, 2007; Thines *et al.*, 2007).

Next to AtKIX8 and AtKIX9, nine other proteins that contain a KIX domain have been described in Arabidopsis (Thakur *et al.*, 2013). Both in plant and non-plant species, such as yeast and humans, the KIX protein family includes HISTONE ACETYLTRANFERASE (HAT) proteins and Mediator subunits (Kumar *et al.*, 2018; Thakur *et al.*, 2013) that are known to function as co-activators through the interaction of their KIX domain with the transactivation domain of transcription factors (Thakur *et al.*, 2014; Kumar *et al.*, 2018). AtKIX8/AtKIX9 and their orthologs, however, are specific to plants and show, except for their N-terminal KIX domain, no similarity to these HAT and Mediator co-activators (Thakur *et al.*, 2013). Instead, they contain an ETHYLENE RESPONSE FACTOR (ERF)-ASSOCIATED AMPHIPHILIC REPRESSION (EAR) motif that allows them to recruit the co-repressor AtTPL (Kagale *et al.*, 2010; Causier *et al.*, 2012; Gonzalez *et al.*, 2015). Through their KIX domain, AtKIX8 and AtKIX9 can simultaneously interact with the transcriptional repressor AtPPD2 (Gonzalez *et al.*, 2015). Hence, AtKIX8/AtKIX9 forms a molecular bridge between AtPPD2 and AtTPL (Supplemental Figure 1A) and, because of that, AtPPD2 can act as a negative transcriptional regulator (Gonzalez *et al.*, 2015).

In Arabidopsis, the activity of the AtPPD-AtKIX repressor complex is regulated by the F-box protein STERILE APETALA (AtSAP) (Wang *et al.*, 2016; Li *et al.*, 2018). Interaction of the repressor complex with AtSAP results in the proteasomal degradation of both AtKIX and AtPPD proteins (Supplemental Figure 1A) (Wang *et al.*, 2016; Li *et al.*, 2018). In accordance with these observations, *AtSAP* overexpression plants produce enlarged rosettes composed of enlarged and dome-shaped leaves (Wang *et al.*, 2016).

Orthologs of the AtPPD, AtKIX, and AtSAP proteins were found in members of both eudicot and monocot species, but appear to be absent from Poaceae species (grasses), suggesting that the PPD-KIX-SAP module was lost in the grass lineage (Gonzalez *et al.*, 2015; Wang *et al.*, 2016). It has been suggested that this might reflect the absence of self-renewing meristemoids in the stomatal lineage of grasses (Liu *et al.*, 2009; Vatén & Bergmann, 2012; Gonzalez *et al.*, 2015; Wang *et al.*, 2016). Several eudicot members, in which orthologs of the *AtPPD* or *AtKIX* genes were mutated or downregulated, including *Medicago truncatula*, soybean (*Glycine max*), blackgram (*Vigna mungo*), and pea (*Pisum sativum*), produced enlarged leaves (Ge *et al.*, 2016; Naito *et al.*, 2017; Kanazashi *et al.*, 2018; Li *et al.*, 2019). Overexpression of *AtSAP* orthologs in *Medicago truncatula*, poplar (*Populus trichocarpa*) and cucumber (*Cucumis sativus* L.) increased leaf size as well (Yordanov *et al.*, 2017; Yang *et al.*, 2018; Yin *et al.*, 2020). Next to enlarged leaves, also increases in the size of other organs, such as stipules, flowers, fruits and seeds, were observed for several of said mutants (Ge *et al.*, 2016; Naito *et al.*, 2017; Kanazashi *et al.*, 2018; Yang *et al.*, 2018; Li *et al.*, 2019; Yin *et al.*, 2020). Control of organ growth by the PPD-KIX repressor complex, together with its post-translational regulation by the F-box protein SAP, thus, seems to be conserved among distinct eudicot species (Schneider et al., 2021). However, differential developmental stages might be targeted by the repressor complex depending on the species. In *Medicago truncatula* plants in which an *AtPPD* ortholog was mutated, for instance, a prolonged period of primary cell division was reported to be responsible for the increase in organ size (Ge *et al.*, 2016), whereas the enlarged leaf phenotype of Arabidopsis *kix8-kix9* and *ami-ppd* mutants was associated with an extended duration of meristemoid division (Gonzalez *et al.*, 2015). The role of conserved regulators can thus vary considerably between different species, illustrating that the translation of knowledge on transcriptional regulators between species is not always straightforward (Nelissen *et al.*, 2014; Kajala *et al.*, 2020). All of the aforementioned eudicot species, in which the function of the PPD-KIX-SAP module was described, belong to the rosid clade. Together with the asterids, the rosids constitute most of the core eudicot species (Supplemental Figure 1B) and their most recent common ancestor existed over 100 million years ago (Wikström *et al.*, 2001). Whether the orthologs of PPD, KIX, and SAP proteins function as regulators of organ growth in asterid members is still unknown.

Here, we report a role for SlKIX8 and SlKIX9 in the regulation of organ growth in the asterid model species tomato (*Solanum lycopersicum*). We used protein interaction assays in yeast to demonstrate that the tomato orthologs of AtKIX8 and AtKIX9 function as SlTPL adaptor proteins for the SlPPD proteins. Next, we used CRISPR-Cas9 genome editing to simultaneously knockout *SlKIX8* and *SlKIX9* in the cultivar Micro-Tom. Double *kix8 kix9* tomato knockout lines produced enlarged, dome-shaped leaves and displayed increased expression of genes orthologous to AtPPD2 target genes. Finally, we demonstrated that *kix8 kix9* and single *kix8* tomato mutants carried larger fruits with increased pericarp thickness, both important agronomic traits for fruit crops, resulting from the production of larger cells.

## Results

### SlKIX8 and SlKIX9 are SlTPL adaptors for SlPPD proteins

To identify the tomato orthologs of the AtKIX8 and AtKIX9 proteins, BLASTP was used. The tomato orthologs of the AtPPD proteins have been described previously (Chini *et al.*, 2017). The SlKIX and SlPPD proteins display a similar domain structure as their Arabidopsis counterparts (Supplemental Figure 2). Amplification of the coding sequences of *SlKIX* and *SlPPD* genes revealed alternative splicing for *SlKIX9*, *SlPPD1*, and *SlPPD2* (Supplemental Figure 2, B–D). Based on an alternative splicing model for *AtKIX9* reported by The Arabidopsis Information Resource (TAIR), we hypothesized that retention of the second *SlKIX9* intron leads to the use of a downstream start codon, generating a splice variant that lacks the N-terminal KIX domain (Supplemental Figure 2B). The splice variants of *SlPPD1* and *SlPPD2* display retention of the Jas intron and part of the Jas intron, respectively, which is located between the two exons encoding the Jas domain (Supplemental Figure 2c,d). These alternative splicing events are proposed to generate premature stop codons (Supplemental Figure 2, C and D), and consequently truncated SlPPD proteins, as was previously shown for *AtPPD1* and *AtPPD2* (Li *et al.*, 2016).

Previously, the interactions between KIX and PPD proteins from Arabidopsis and pea were analyzed *in vivo* and in detail in yeast (Gonzalez *et al.*, 2015; Li *et al.*, 2019). To determine whether SlKIX8, SlKIX9, SlPPD1, and SlPPD2 are part of a similar protein complex, we performed yeast two-hybrid (Y2H) assays. For these assays, the splice variants with the most complete coding sequence (shown in Supplemental Figure 2) were used. In case of the SlKIX proteins, these possessed the KIX domain, which was shown to be essential for mediating the interaction between KIX and PPD proteins from Arabidopsis and pea (Gonzalez *et al.*, 2015; Li *et al.*, 2019). Direct interaction between the SlKIX and SlPPD proteins could be observed (Figure 1A). Next, we evaluated whether the SlKIX proteins were able to interact with SlTPL1 (Figure 1B), which is the most closely related tomato ortholog of the Arabidopsis co-repressor AtTPL (Hao *et al.*, 2014). As only SlKIX8 was capable of interacting with SlTPL1 in the Y2H assays (Figure 1B), we also assessed the interaction between the SlKIX proteins and the five additional SlTPL proteins that were reported in tomato (Figure 1B). In addition to SlTPL1, SlKIX8 also interacted with SlTPL2, SlTPL4, and SlTPL6, whereas SlKIX9 could solely interact with SlTPL2 (Figure 1B). By means of yeast three-hybrid (Y3H) assays, we subsequently demonstrated that the SlKIX proteins can form a molecular bridge between these SlTPL proteins and the SlPPD proteins (Figure 1, C and D). In Arabidopsis, both AtKIX8 and AtKIX9 were reported to interact with the F-box protein AtSAP, resulting in their post-translational degradation (Li *et al.*, 2018). However, we only observed interaction between SlKIX8 and the tomato ortholog of AtSAP (Figure 1E). Taken together, our results suggest that in tomato, SlKIX8 and SlKIX9 function as SlTPL adaptors for the SlPPD proteins, similar to their orthologs in Arabidopsis.

**Figure 1.**
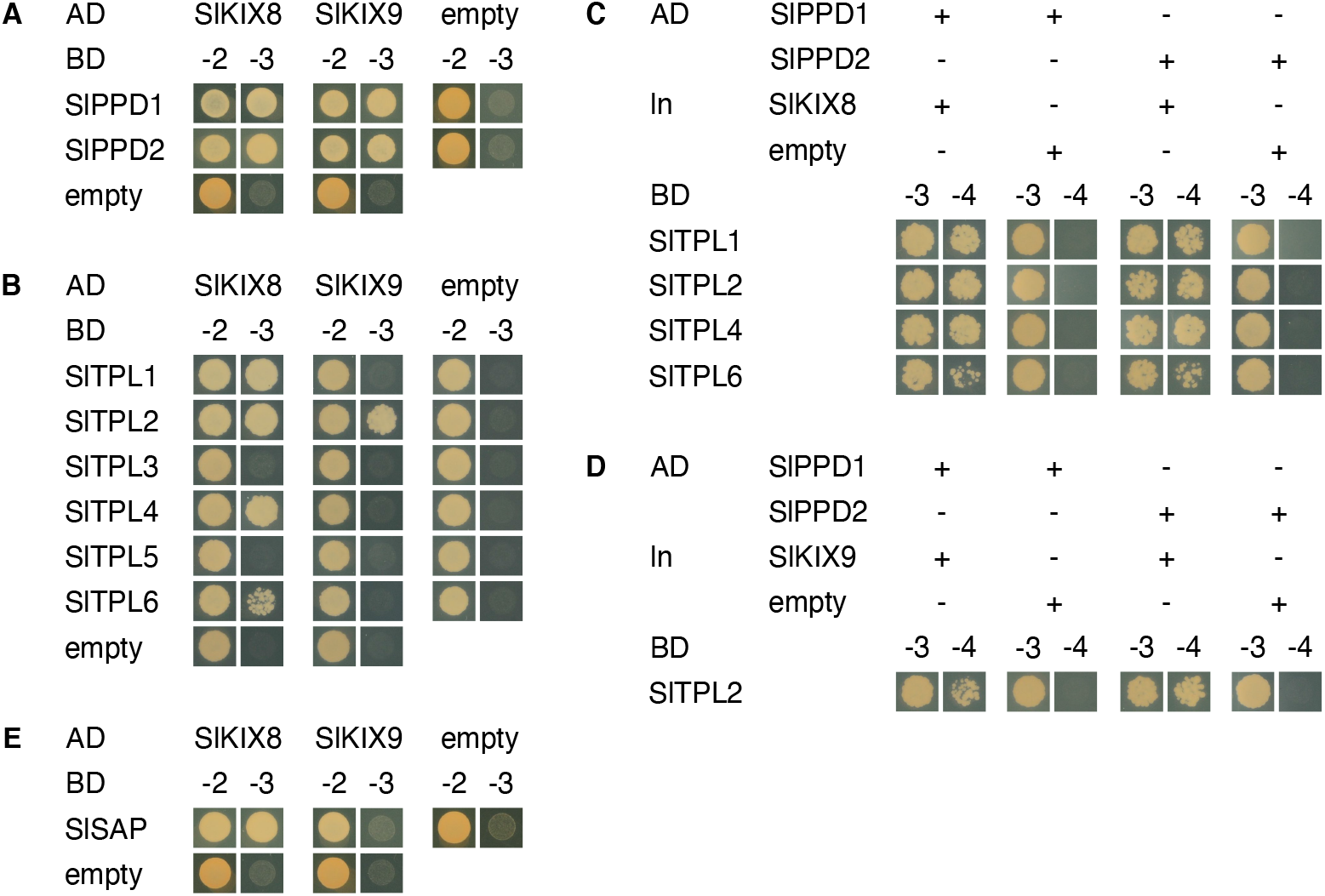
SlKIX8 and SlKIX9 are SlTPL adaptors for SlPPD proteins. (A−B) Y2H interaction analysis of SlKIX8 and SlKIX9 with SlPPD (A) and SlTPL (B) proteins. Yeast transformants expressing bait (BD) and prey (AD) proteins were dropped on control medium lacking Leu and Trp (−2) or selective medium additionally lacking His (−3). (C−D) Y3H interaction analysis to test the bridging capacity of SlKIX8 (c) and SlKIX9 (d) to mediate the SlPPD-SlTPL interaction. Yeast transformants expressing bait (BD), bridge (ln), and prey (AD) proteins were dropped on control medium lacking Leu, Trp, and Ura (−3) or selective medium additionally lacking His (−4). (E) Y2H interaction analysis of SlKIX proteins with SlSAP. Yeast transformants expressing bait (BD) and prey (AD) proteins were dropped on control medium lacking Leu and Trp (−2) or selective medium additionally lacking His (−3). Empty vectors were used in all control assays. Abbreviations: AD, activation domain; BD, binding domain; ln, linker.

### CRISPR-Cas9 genome editing of *SlKIX8* and *SlKIX9* leads to enlarged dome-shaped leaves

To investigate the *in planta* role of SlKIX8 and SlKIX9, double *kix8 kix9* loss-of-function mutants (cultivar Micro-Tom) were generated using Clustered Regularly Interspaced Short Palindromic Repeats-CRISPR associated protein 9 (CRISPR-Cas9) genome editing (Figure 2A). A rippled, dome-shaped leaf phenotype could already be observed in regenerated double *kix8 kix9* tomato knockout (T0) plants (Supplemental Figure 3). Likewise, the progeny of two independent T1 plants mono- or biallelic for out-of-frame mutations at both the *SlKIX8* and *SlKIX9* loci (Supplemental Figure 4, A and B), displayed dome-shaped leaves with uneven leaf laminae (Figure 2, B and C). The main shoot length of these double *kix8 kix9* mutants was reduced compared with that of wild-type plants (Supplemental Table 1). Single *kix8* mutants (Supplemental Figure 4, A and B), obtained by pollinating the *kix8 kix9*^#1^ (T1) line with wild-type pollen, exhibited an intermediate leaf phenotype (Figure 2, B and C), whereas single *kix9* plants (Supplemental Figure 4, A and B) did not show any visible phenotype (Figure 2, B and C) as was noted for Arabidopsis *kix8* and *kix9* single mutants (Gonzalez *et al.*, 2015). These observations suggest that SlKIX8 and SlKIX9 may have partially redundant roles in regulating tomato leaf growth.

**Figure 2.**
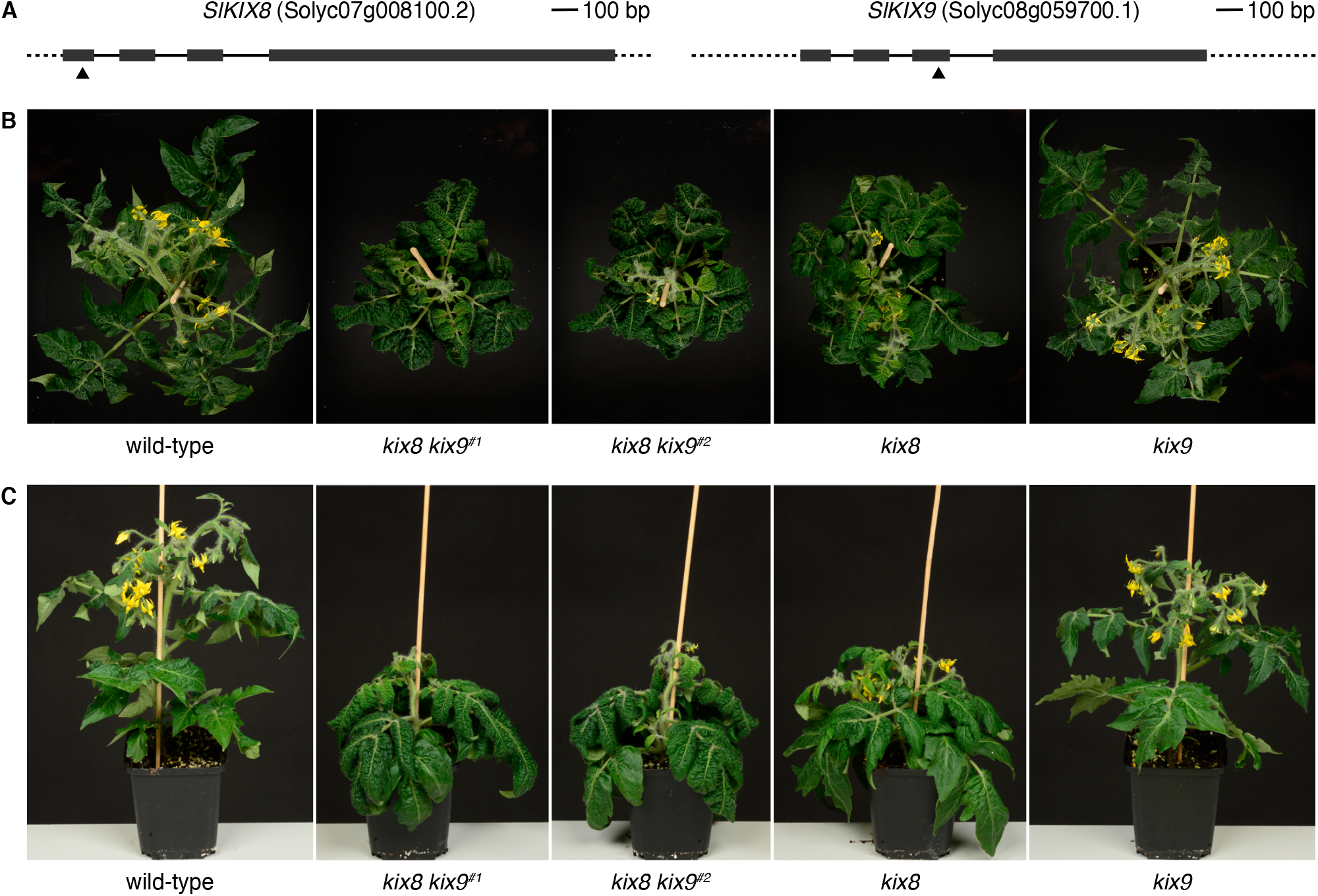
CRISPR-Cas9 genome editing of tomato *SlKIX8* and *SlKIX9* causes a rippled, dome-shaped leaf phenotype. (A) Schematic representation of *SlKIX8* and *SlKIX9* with location of the CRISPR-Cas9 cleavage sites. Dark grey boxes represent exons. Cas9 cleavage sites for guide RNAs are indicated with arrowheads. (B−C) Representative wild-type, *kix8 kix9*^#1^, *kix8 kix9^#2^*, *kix8*, and *kix9* plants grown in soil for 1 month under 16:8 photoperiods with daytime and nighttime temperatures of 26–29°C and 18–20°C, respectively, were photographed from the top (B) and the front (C).

Given that the leaf shape phenotype was most pronounced for double *kix8 kix9* tomato mutants, phenotypical analyses were performed on leaf eight and compared with those of the corresponding wild-type leaf. First, leaf fresh weight was determined, which was approximately 30% higher for *kix8 kix9* leaves compared with wild-type leaves (Figure 3A). Likewise, the fresh weight of the terminal leaflets of these *kix8 kix9* leaves was increased with approximately 40% compared with those of wild-type leaves (Figure 3B). Next, the area of terminal leaflets was measured before (projected area) and after (real area) terminal leaflets were cut to flatten them (Figure 3, C and D) (Baekelandt *et al.*, 2018). After flattening the terminal leaflets, those of *kix8 kix9* mutants displayed an area that was approximately 40% larger than corresponding wild-type leaflets (Figure 3E). In addition, the decrease in projected-to-real terminal leaflet area was about two times bigger for *kix8 kix9* plants compared with wild-type plants (Figure 3F), demonstrating the alteration in *kix8 kix9* leaflet shape. These measurements, thus, substantiate the enlarged, dome-shaped leaf phenotype of double *kix8 kix9* tomato knockout plants.

**Figure 3.**
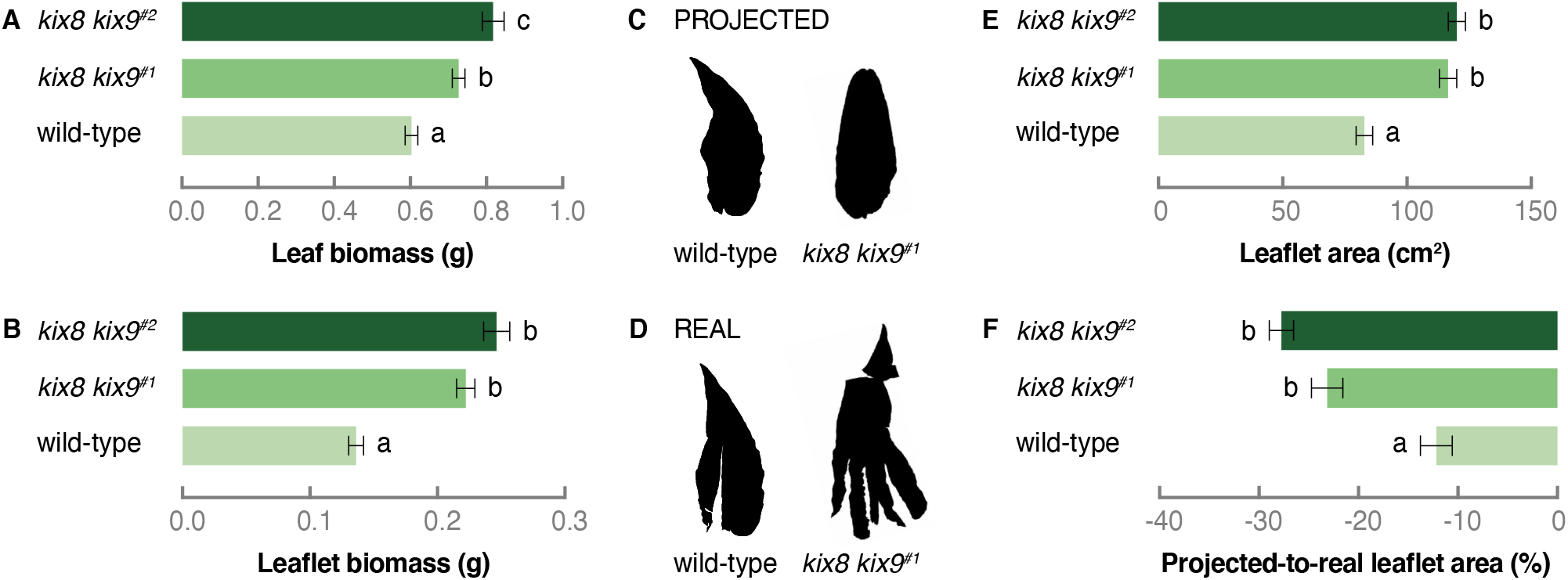
Tomato *kix8 kix9* plants produce enlarged, dome-shaped leaves. (A−B) Biomass of leaf eight (from the top) (A) and its terminal leaflet (B). The eighth leaf (from the top) was harvested from plants grown in soil for 2 months under 16:8 photoperiods with daytime and nighttime temperatures of 26–29°C and 18–20°C, respectively. Bars represent mean biomass relative to the mean of wild-type biomass values. Error bars denote standard error (n = 31–40). Statistical significance was determined by ANOVA followed by Tukey post-hoc analysis (P < 0.05; indicated by different letters). (C−D) The terminal leaflet area was measured before (projected, C) and after (real, D) the terminal leaflet of the eighth leaf was cut to flatten it. (E−F) Area (E) and projected-to-real area (F) of the terminal leaflet of the eighth leaf. Bars represent mean area relative to the mean of wild-type area values. Error bars denote standard error (n = 31–40). Statistical significance was determined by ANOVA followed by Tukey post-hoc analysis (P < 0.05; indicated by different letters).

### Orthologs of AtPPD2 target genes are upregulated in leaves of tomato *kix8 kix9* mutants

To gain further insight into the function of SlKIX8 and SlKIX9 in tomato plants, we made use of public transcriptome data (cultivar Micro-Tom) (Zouine *et al.*, 2017) to investigate the gene expression patterns of *SlKIX8*, *SlKIX9*, *SlPPD1*, and *SlPPD2* in different tissues and throughout distinct developmental stages. A survey of these publicly available transcriptome data revealed that *SlKIX8* was low expressed in all examined tissues and that *SlKIX9* expression was (almost) absent in most tissues (Figure 4A and Supplemental Table 2). In all investigated tissues, the transcript level of *SlPPD2* was higher than that of *SlPPD1* (Figure 4A and Supplemental Table 2). Next, we looked up the gene expression patterns of the putative tomato orthologs of Arabidopsis *DWARF IN LIGHT 1* (*AtDFL1*), *AT-HOOK MOTIF CONTAINING NUCLEAR LOCALISED PROTEIN 17* (*AtAHL17*), and *SCHLAFMUTZE* (*AtSMZ*), which were top-ranked in the list of differentially expressed genes in Arabidopsis *ami-ppd* leaves and strongly upregulated in Arabidopsis *kix8-kix9* leaves (Gonzalez *et al.*, 2015). Expression of all three tomato genes, *SlDFL1*, *SlAHL17*, and *APETALA 2d* (*SlAP2d*), was confirmed in tomato leaves (Figure 4B and Supplemental Table 2). To verify the potential differential expression of these genes in tomato *kix8 kix9* mutants compared with wild-type plants, we performed a quantitative real-time PCR (qPCR) analysis on the terminal leaflet of growing leaves and found that the transcription of all three genes was upregulated in *kix8 kix9* mutants (Figure 4C). Furthermore, the expression of *SlKIX8* and *SlKIX9* was increased in *kix8 kix9* plants compared with wild-type plants (Figure 4D), suggesting negative feedback of the SlPPD-SlKIX complex on the expression of *SlKIX8* and *SlKIX9*. These findings indicate that SlKIX8 and SlKIX9 are required for the repression of tomato genes orthologous to three AtPPD2 target genes.

**Figure 4.**
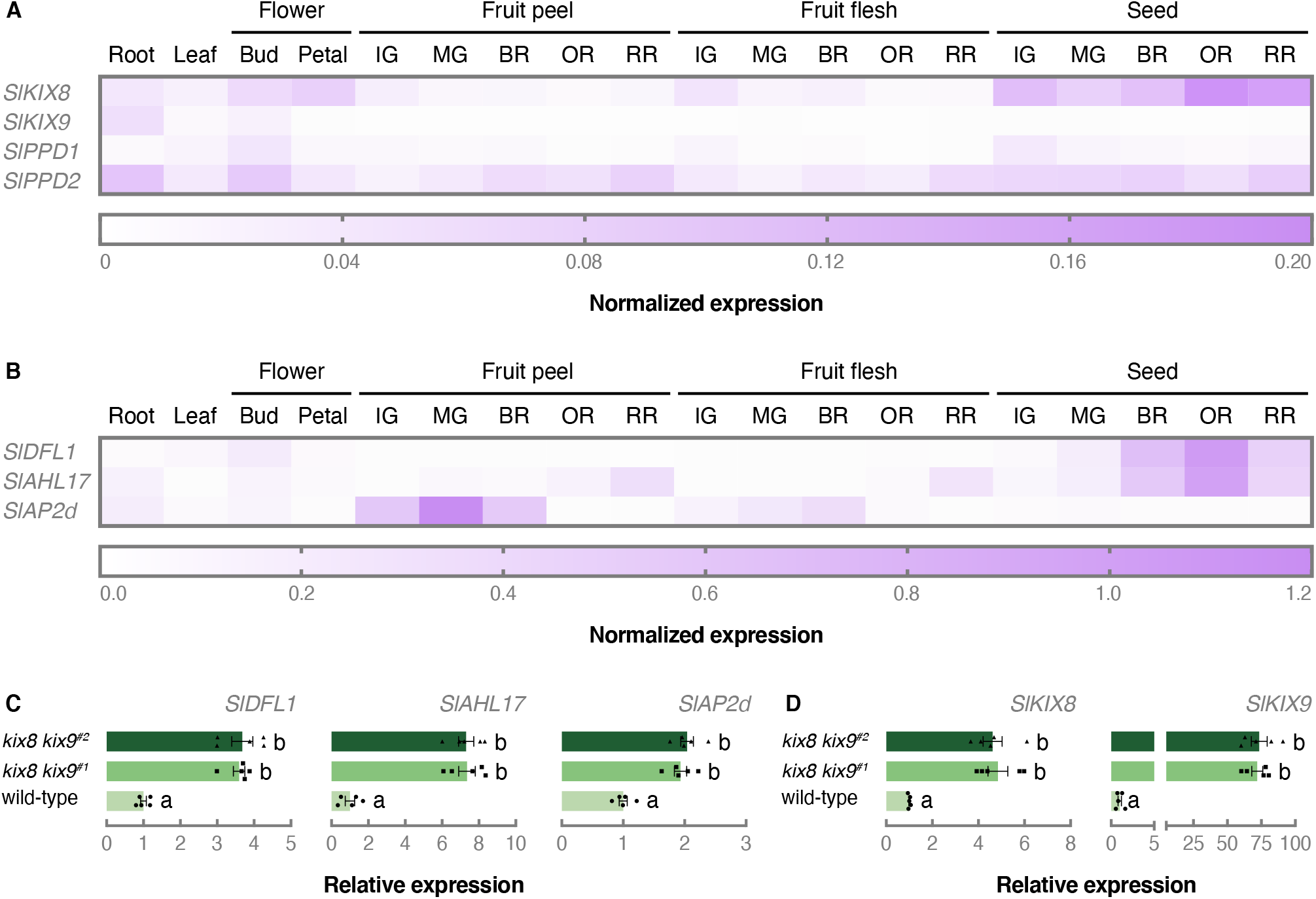
SlKIX8 and SlKIX9 are required for the repression of putative SlPPD target genes. (A−B) Normalized expression profiles of *SlKIX8*, *SlKIX9*, *SlPPD1*, *SlPPD2* (A), *SlDFL1*, *SlAHL17*, and *SlAP2d* (B) in different tomato organs and developmental stages (cultivar Micro-Tom). Expression data was obtained from TomExpress (Zouine *et al.*, 2017) and can be found in Supplemental Table 1. (C−D) Relative expression of *SlDFL1*, *SlAHL17*, *SlAP2d* (C), *SlKIX8*, and *SlKIX9* (D) in terminal leaflets of not fully developed leaves analyzed by qPCR. The terminal leaflet from the second leaf (from the top) was harvested from plants grown in soil for 3 weeks under 16:8 photoperiods with daytime and nighttime temperatures of 26–29°C and 18–20°C, respectively. Bars represent mean expression relative to the mean of wild-type expression values. Error bars denote standard error (n = 5). Individual wild-type (•), *kix8 kix9*^#1^ (▪), and *kix8 kix9*^#2^ (▴) values are shown. Statistical significance was determined by ANOVA followed by Tukey post-hoc analysis (P < 0.05; indicated by different letters). Abbreviations: IG, immature green; MG, mature green; BR, breaker; OR, orange; RR, red ripe.

### Tomato *kix8 kix9* mutants produce enlarged fruits due to increased cell expansion

In multiple eudicot species that belong to the rosid order of Fabales, orthologs of the KIX and PPD proteins have been reported to negatively regulate seed pod size (Ge *et al.*, 2016; Naito *et al.*, 2017; Kanazashi *et al.*, 2018; Li *et al.*, 2019). Moreover, the cucumber ortholog of the F-box protein AtSAP was shown to positively regulate fruit size (Yang *et al.*, 2018). To examine whether the SlKIX proteins might also have a role in determining fruit size in the asterid model species tomato, we investigated if the development of reproductive organs was affected in tomato plants in which *SlKIX8* and/or *SlKIX9* function was disturbed.

Fruits that developed on inflorescences of the main shoot were harvested from each genotype when the ratio of ripe to unripe fruits was 65–85%, since we noted a significant delay in flowering time for *kix8 kix9* mutants compared with wild-type plants (Supplemental Figure 5). The fresh weight of individual ripe tomatoes produced by *kix8 kix9* and *kix8* plants was increased by approximately 15% and 30%, respectively, compared with those produced by wild-type plants (Table 1, Figure 5A). Cutting along the equatorial plane revealed that *kix8 kix9* and *kix8* fruits displayed an approximate increase of 50% in pericarp thickness compared with wild-type fruits, while no change was observed for *kix9* fruits (Table 1, Figure 5B). Red fruit total biomass per plant was increased for *kix8* and *kix9* but not *kix8 kix9* mutants compared with wild-type plants (Table 1). We noted that the number of red fruits per plant remained similar for *kix8* plants and increased for *kix9* plants while it was lower for *kix8 kix9* plants than for wild-type plants (Table 1), the latter possibly resulting from an increase in flower abortion ratio within inflorescences (Table 1). To better estimate the effect of *SlKIX* loss-of-function on fruit yield, we performed a second experiment for which we harvested fruits not only from inflorescences on the main shoot but also on the axillary shoots from *kix8 kix9* and wild-type plants. Whereas the number of inflorescences on the main shoot was unaffected in *kix8 kix9* mutants compared to wild-type plants, the number of inflorescences on the axillary shoots was reduced by approximately 50% (Supplemental Table 3), indicating a delay in axillary fruit development. Although the higher biomass of *kix8 kix9* fruits accompanied by increased pericarp thickness compared with wild-type fruits was confirmed (Supplemental Table 3), the reduced axillary branching reduced total fruit yield per plant by approximately 30% (Supplemental Table 3). In addition, we observed that *kix8 kix9* fruits contained approximately 45% less seeds than wild-type fruits, though seed size was unaffected (Supplemental Table 3).

**Figure 5.**
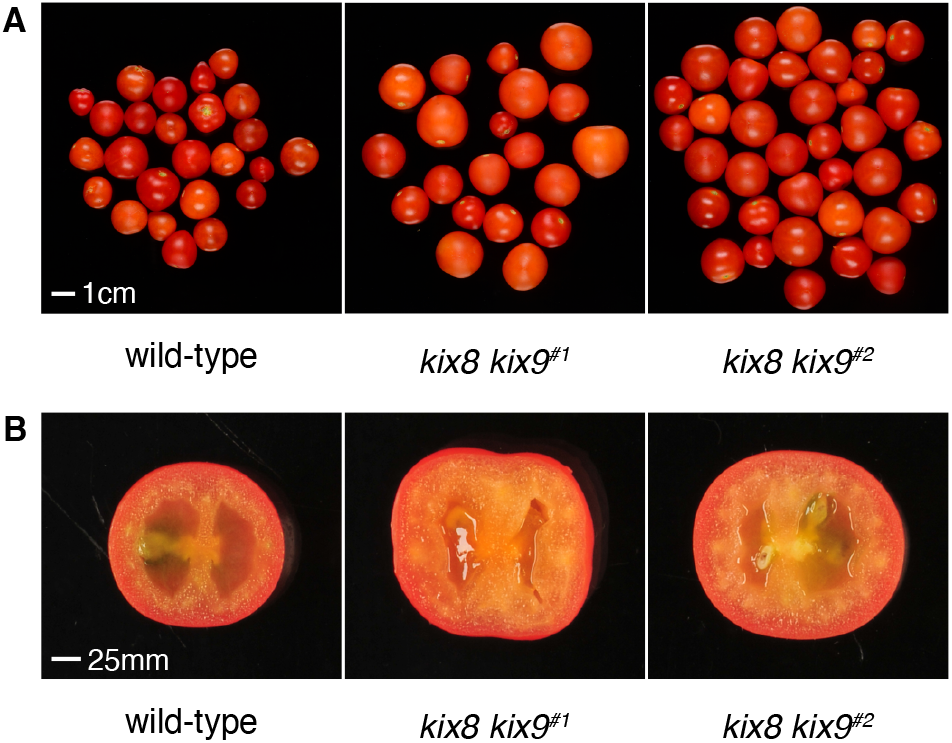
Tomato *kix8 kix9* plants produce enlarged fruits that display increased pericarp thickness. Representative red ripe fruits produced by wild-type, *kix8 kix9*^#1^, and *kix8 kix9*^#2^ plants. (B) Equatorial sections of representative red ripe fruits produced by wild-type, *kix8 kix9*^#1^, and *kix8 kix9^#2^* plants. Plants were grown in soil under 16:8 photoperiods with daytime and nighttime temperatures of 26–29°C and 18–20°C, respectively.

**Table 1.** Tomato *kix8* and *kix8 kix9* plants produce bigger fruits with increased pericarp thickness.

|  | wild-type | *kix8* | *kix9* | *kix8 kix9*<sup>#1</sup> | *kix8 kix9*<sup>#2</sup> |
| --- | --- | --- | --- | --- | --- |
| **Inflorescence parameters:** |  |  |  |  |  |
| Number of inflorescences | 4.00 ± 0.17<sup>abc</sup> | 4.44 ± 0.24<sup>ab</sup> | 4.77 ± 0.22<sup>b</sup> | 3.78 ± 0.15<sup>ac</sup> | 3.44 ± 0.18<sup>c</sup> |
| Number of flowers per inflorescence | 6.15 ± 0.67<sup>ab</sup> | 6.01 ± 0.33<sup>ab</sup> | 6.74 ± 0.42<sup>a</sup> | 5.77 ± 0.38<sup>b</sup> | 5.53 ± 0.32<sup>b</sup> |
| Number of pollinated flowers per inflorescence | 5.70 ± 0.70<sup>a</sup> | 4.13 ± 0.66<sup>b</sup> | 6.67 ± 0.41<sup>a</sup> | 4.53 ± 0.55<sup>b</sup> | 4.48 ± 0.55<sup>b</sup> |
| Pollinated/total number of flowers per inflorescence (%) | 93.13 ± 3.65<sup>ac</sup> | 69.03 ± 10.01<sup>b</sup> | 99.04 ± 1.21<sup>c</sup> | 77.63 ± 7.49<sup>bd</sup> | 80.96 ± 8.12<sup>ad</sup> |
| **Fruit parameters:** |  |  |  |  |  |
| Red fruit biomass (g) | 4.40 ± 0.65<sup>ab</sup> | 5.78 ± 0.57<sup>c</sup> | 3.63 ± 0.41<sup>b</sup> | 5.00 ± 0.56<sup>ac</sup> | 5.12 ± 0.59<sup>ac</sup> |
| Red fruit pericarp thickness (mm) | 2.67 ± 0.08<sup>a</sup> | 4.08 ± 0.16<sup>b</sup> | 2.65 ± 0.13<sup>a</sup> | 3.97 ± 0.08<sup>b</sup> | 4.16 ± 0.13<sup>b</sup> |
| Red fruit pericarp thickness/radius (%) | 21.44 ± 0.67<sup>a</sup> | 31.78 ± 1.31<sup>b</sup> | 22.59 ± 0.85<sup>a</sup> | 32.07 ± 0.84<sup>b</sup> | 32.28 ± 0.90<sup>b</sup> |
| **Fruit yield parameters:** |  |  |  |  |  |
| Number of red fruits | 16.56 ± 1.97<sup>a</sup> | 15.44 ± 0.97<sup>a</sup> | 26.67 ± 1.96<sup>b</sup> | 13.56 ± 0.60<sup>a</sup> | 12.44 ± 0.75<sup>a</sup> |
| Red fruit yield (g) | 69.85 ± 6.25<sup>ac</sup> | 88.71 ± 4.64<sup>ab</sup> | 97.00 ± 6.97<sup>b</sup> | 67.09 ± 2.18<sup>c</sup> | 62.51 ± 3.63<sup>c</sup> |
| Plants were grown in soil for 3.5–4.5 months under 15:9 photoperiods with daytime and nighttime temperatures of 25–29°C and 17–19°C, respectively. Inflorescence parameters were measured for the main shoot and red fruits that developed on these inflorescences were harvested from each genotype when the ratio of ripe to unripe fruits was 65–85%. Data are mean ± standard error (n = 7–9). Statistical significance was determined by ANOVA followed by Tukey post-hoc analysis (P < 0.05; indicated by different letters). |  |  |  |  |  |

To determine the influence of an altered sink–source relationship between *kix8 kix9*, *kix8*, *kix9*, and wild-type plants, a follow-up experiment was performed in which fruit production was restricted. To do so, only the first two inflorescences on the main shoot, each carrying a maximum of six fruits, were kept per plant and growth parameters were documented from ovary stage until red ripe stage. Even though *kix8 kix9* and *kix8* mutants carried ovaries that were approximately 35% smaller than wild-type plants (Table 2), final red fruit biomass was unaffected (Table 2). Moreover, the increase in pericarp thickness was confirmed in these restricted conditions for *kix8 kix9* and *kix8* fruits from 15 days post anthesis (DPA) onwards, with ripe *kix8 kix9* and *kix8* fruits harboring a pericarp that was approximately 20% thicker than wild-type fruits (Table 2). To explore the cellular cause of this change in pericarp size, the number of cell layers and cell sizes were quantified. At 30 DPA, the number of cell layers across the pericarp was similar in wild-type, *kix8*, *kix9*, and *kix8 kix9* fruits (Figure 6A). The average pericarp cell area, however, was increased in all mutant fruits compared with wild-type fruits (Figure 6, B and C). The increased cell area resulted from the appearance of very large cells within the pericarp of *kix8*, *kix9*, and *kix8 kix9* and a decreased proportion of the smallest cells in *kix8* and *kix8 kix9* compared to wild-type (Figure 6D). Altogether, these data demonstrate that knocking out *SlKIX8* on its own or *SlKIX8* together with *SlKIX9* results in the production of enlarged tomato fruits with increased pericarp thickness, suggesting that SlKIX8 and/or SlKIX9 are involved in the regulation of tomato fruit growth.

**Table 2.**
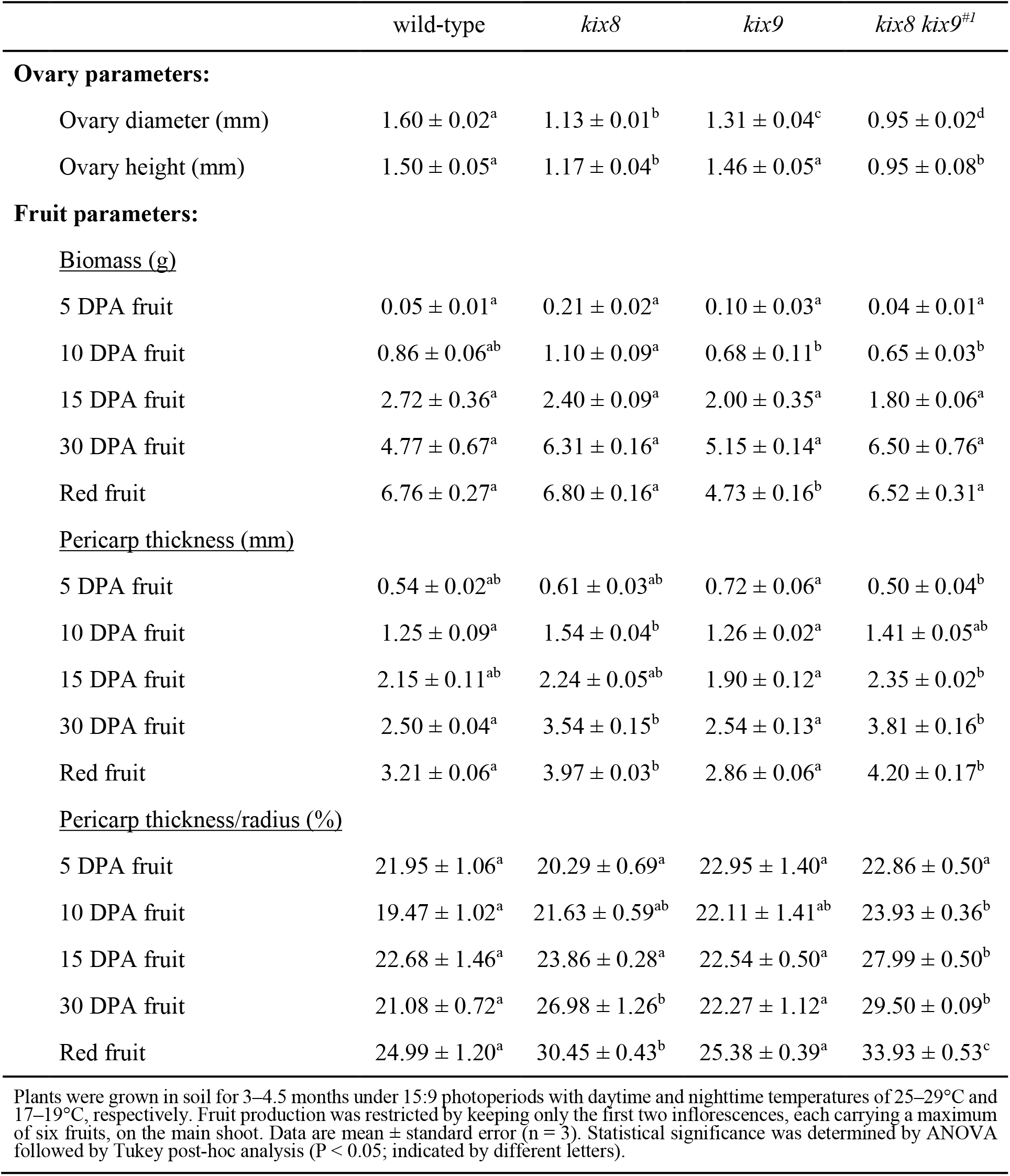
Pericarp thickness of fruits from tomato *kix8* and *kix8 kix9* plants grown in restricted conditions is increased.

|  | wild-type | *kix8* | *kix9* | *kix8 kix9*<sup>#1</sup> |
| --- | --- | --- | --- | --- |
| **Ovary parameters:** |  |  |  |  |
| Ovary diameter (mm) | 1.60 ± 0.02<sup>a</sup> | 1.13 ± 0.01<sup>b</sup> | 1.31 ± 0.04<sup>c</sup> | 0.95 ± 0.02<sup>d</sup> |
| Ovary height (mm) | 1.50 ± 0.05<sup>a</sup> | 1.17 ± 0.04<sup>b</sup> | 1.46 ± 0.05<sup>a</sup> | 0.95 ± 0.08<sup>b</sup> |
| **Fruit parameters:** |  |  |  |  |
| <u>Biomass (g)</u> |  |  |  |  |
| 5 DPA fruit | 0.05 ± 0.01<sup>a</sup> | 0.21 ± 0.02<sup>a</sup> | 0.10 ± 0.03<sup>a</sup> | 0.04 ± 0.01<sup>a</sup> |
| 10 DPA fruit | 0.86 ± 0.06<sup>ab</sup> | 1.10 ± 0.09<sup>a</sup> | 0.68 ± 0.11<sup>b</sup> | 0.65 ± 0.03<sup>b</sup> |
| 15 DPA fruit | 2.72 ± 0.36<sup>a</sup> | 2.40 ± 0.09<sup>a</sup> | 2.00 ± 0.35<sup>a</sup> | 1.80 ± 0.06<sup>a</sup> |
| 30 DPA fruit | 4.77 ± 0.67<sup>a</sup> | 6.31 ± 0.16<sup>a</sup> | 5.15 ± 0.14<sup>a</sup> | 6.50 ± 0.76<sup>a</sup> |
| Red fruit | 6.76 ± 0.27<sup>a</sup> | 6.80 ± 0.16<sup>a</sup> | 4.73 ± 0.16<sup>b</sup> | 6.52 ± 0.31<sup>a</sup> |
| <u>Pericarp thickness (mm)</u> |  |  |  |  |
| 5 DPA fruit | 0.54 ± 0.02<sup>ab</sup> | 0.61 ± 0.03<sup>ab</sup> | 0.72 ± 0.06<sup>a</sup> | 0.50 ± 0.04<sup>b</sup> |
| 10 DPA fruit | 1.25 ± 0.09<sup>a</sup> | 1.54 ± 0.04<sup>b</sup> | 1.26 ± 0.02<sup>a</sup> | 1.41 ± 0.05<sup>ab</sup> |
| 15 DPA fruit | 2.15 ± 0.11<sup>ab</sup> | 2.24 ± 0.05<sup>ab</sup> | 1.90 ± 0.12<sup>a</sup> | 2.35 ± 0.02<sup>b</sup> |
| 30 DPA fruit | 2.50 ± 0.04<sup>a</sup> | 3.54 ± 0.15<sup>b</sup> | 2.54 ± 0.13<sup>a</sup> | 3.81 ± 0.16<sup>b</sup> |
| Red fruit | 3.21 ± 0.06<sup>a</sup> | 3.97 ± 0.03<sup>b</sup> | 2.86 ± 0.06<sup>a</sup> | 4.20 ± 0.17<sup>b</sup> |
| <u>Pericarp thickness/radius (%)</u> |  |  |  |  |
| 5 DPA fruit | 21.95 ± 1.06<sup>a</sup> | 20.29 ± 0.69<sup>a</sup> | 22.95 ± 1.40<sup>a</sup> | 22.86 ± 0.50<sup>a</sup> |
| 10 DPA fruit | 19.47 ± 1.02<sup>a</sup> | 21.63 ± 0.59<sup>ab</sup> | 22.11 ± 1.41<sup>ab</sup> | 23.93 ± 0.36<sup>b</sup> |
| 15 DPA fruit | 22.68 ± 1.46<sup>a</sup> | 23.86 ± 0.28<sup>a</sup> | 22.54 ± 0.50<sup>a</sup> | 27.99 ± 0.50<sup>b</sup> |
| 30 DPA fruit | 21.08 ± 0.72<sup>a</sup> | 26.98 ± 1.26<sup>b</sup> | 22.27 ± 1.12<sup>a</sup> | 29.50 ± 0.09<sup>b</sup> |
| Red fruit | 24.99 ± 1.20<sup>a</sup> | 30.45 ± 0.43<sup>b</sup> | 25.38 ± 0.39<sup>a</sup> | 33.93 ± 0.53<sup>c</sup> |

Plants were grown in soil for 3–4.5 months under 15:9 photoperiods with daytime and nighttime temperatures of 25–29°C and 17–19°C, respectively. Fruit production was restricted by keeping only the first two inflorescences, each carrying a maximum of six fruits, on the main shoot. Data are mean ± standard error (n = 3). Statistical significance was determined by ANOVA followed by Tukey post-hoc analysis (P < 0.05; indicated by different letters).

**Figure 6.**
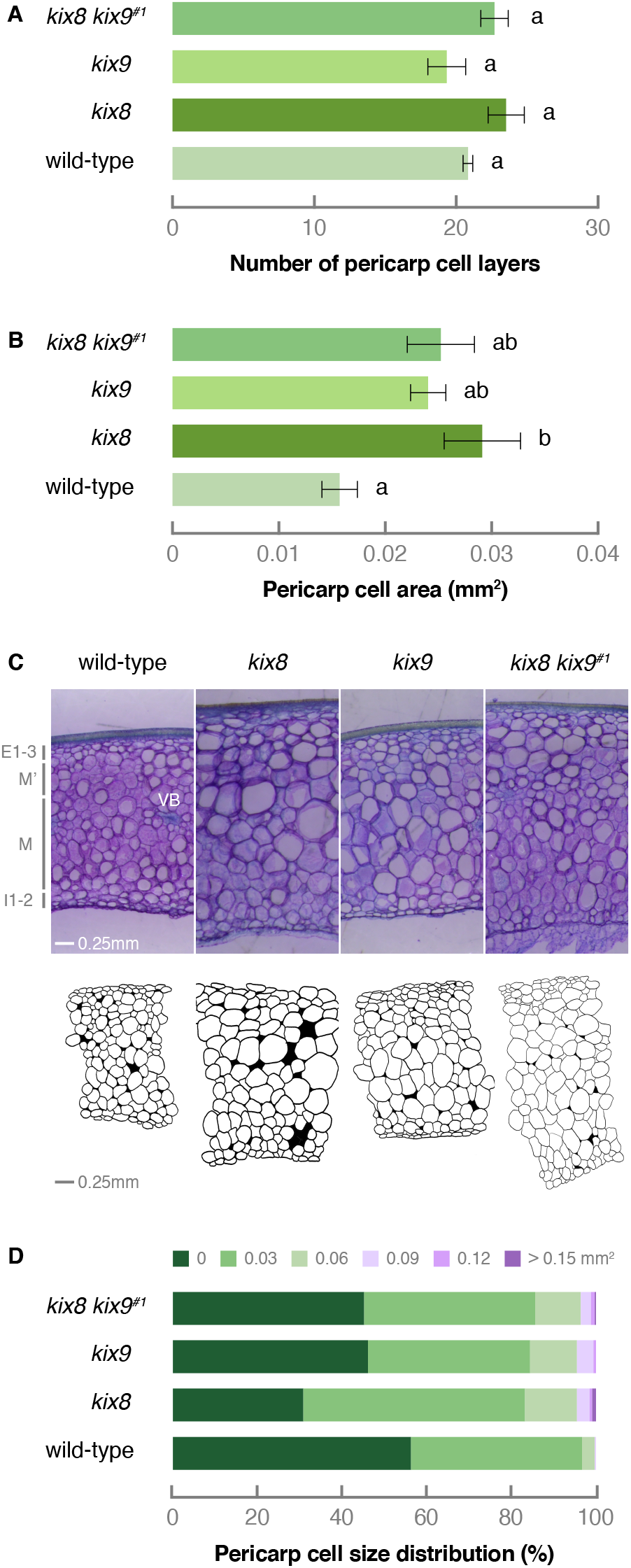
Increased pericarp thickness of *kix8* and *kix8 kix9* fruits results from the production of larger cells.(A) Number of pericarp cell layers in wild-type, *kix8*, *kix9*, and *kix8 kix9*^#1^ fruit. (B) Pericarp cell area in wild-type, *kix8*, *kix9*, and *kix8 kix9*^#1^ fruit. (C) Microtom pericarp sections and drawings of a representative wild-type, *kix8*, *kix9*, and *kix8 kix9*^#1^ fruit. (D) Pericarp cell size distribution in wild-type, *kix8*, *kix9*, and *kix8 kix9*^#1^ fruit. Plants were grown in soil under 15:9 photoperiods with daytime and nighttime temperatures of 25–29°C and 17–19°C, respectively. Fruit production was restricted by keeping only the first two inflorescences, each carrying a maximum of six fruits, on the main shoot. Fruit was harvested at 30 days post anthesis. Data are mean ± standard error (n = 2−4). Statistical significance was determined by ANOVA followed by Tukey post-hoc analysis (P < 0.05; indicated by different letters). Abbreviations: E, outer epidermis layer; M, mesocarp layer; I, inner epidermis layer; VB, vascular bundle.

## Discussion

### KIX8 and KIX9 are regulators of leaf growth in distinct eudicot species

In Arabidopsis, the asymmetric cell division of meristemoids and leaf growth positively are restricted by a transcriptional repressor complex in which the co-repressor AtTPL is recruited to AtPPD2 by AtKIX8/AtKIX9 (White, 2006; Gonzalez *et al.*, 2015). Members of this repressor complex were shown to regulate leaf size and shape in a variety of species that belong to different orders of the rosids (Gonzalez *et al.*, 2015; Ge *et al.*, 2016; Naito *et al.*, 2017; Kanazashi *et al.*, 2018; Li *et al.*, 2019), suggesting that the repressor complex is a conserved regulator of leaf growth among rosid eudicots (Schneider et al., 2021). Here, we demonstrate that the tomato orthologs of AtKIX8 and AtKIX9 act as SlTPL adaptors for SlPPD proteins and, thereby, regulate leaf growth in tomato plants. Tomato is a model species of the asterid clade that also includes tobacco (*Nicotiana tabacum*), carrot (*Daucus carota*) and sunflower (*Helianthus annuus*). In the rosid species Arabidopsis and pea, the interaction between KIX and PPD proteins is described to occur through the N-terminal KIX and PPD domain, respectively (Gonzalez *et al.*, 2015; Li *et al.*, 2019). This is likely to be the case in tomato as well, in which the SlKIX and SlPPD proteins display a similar domain structure. The interaction between tomato SlKIX8/SlKIX9 and the SlTPL co-repressors is expected to occur via the EAR motif present in the SlKIX proteins (Kagale *et al.*, 2010; Causier *et al.*, 2012), as was shown for their Arabidopsis and pea orthologs (Gonzalez *et al.*, 2015; Li *et al.*, 2019).

Tomato *kix8 kix9* plants exhibited an enlarged, dome-shaped leaf phenotype, similar to the phenotype observed in Arabidopsis *kix8-kix9* and *ami-ppd* mutants (Gonzalez *et al.*, 2015). Moreover, terminal leaflets of young tomato *kix8 kix9* leaves displayed an increased expression of three putative SlPPD target genes, *SlDFL1*, *SlAHL17*, and *SlAP2d*, of which the orthologs were strongly upregulated in Arabidopsis *kix8-kix9* and *ami-ppd* leaves (Gonzalez *et al.*, 2015). Therefore, the SlPPD-SlKIX complex might control similar molecular processes during leaf development as in Arabidopsis (White, 2006; Gonzalez *et al.*, 2015). Taken together, we can conclude that both in rosid and asterid species, KIX8 and/or KIX9 assist PPD proteins in repressing distinct downstream target genes and in regulating leaf size and shape.

### Partial redundancy of SlKIX8 and SlKIX9

In Arabidopsis, AtKIX8 and AtKIX9 were reported to have partially redundant roles in controlling leaf growth (Gonzalez *et al.*, 2015). The intermediate and absent leaf phenotype of tomato single *kix8* and *kix9* mutants, respectively, compared with the markedly enlarged, dome-shaped leaf phenotype of tomato *kix8 kix9* plants, suggests partial redundancy of SlKIX8 and SlKIX9 in tomato leaf development as well. In fruits, we observed stronger phenotypes for single *kix8* than *kix9* mutants. In line with these phenotypes, *SlKIX9* expression is (almost) absent in most tomato tissues, whereas *SlKIX8* displays an overall higher expression level in the examined tissues. In tomato *kix8 kix9* tomato leaflets, however, the transcript levels of not only *SlKIX8* but also *SlKIX9* were increased compared with wild-type leaflets, suggesting negative feedback of the SlPPD-SlKIX complex on the expression of both *SlKIX8* and *SlKIX9*. In yeast cells, interaction with SlTPL2 was observed for both SlKIX8 and SlKIX9, but SlKIX8 could additionally interact with SlTPL1, SlTPL4, SlTPL6, and SlSAP. A previous study showed that from the six SlTPL genes, *SlTPL1* had the highest overall expression in the examined tissues and developmental stages, while *SlTPL2* was expressed to a much lesser extent (Hao *et al.*, 2014). The expression of *SlTPL4* dominated in ripening fruit and *SlTPL6* transcripts were almost absent in all investigated tissues (Hao *et al.*, 2014). Furthermore, SlTPL6 was suggested to have lost its functionality (Hao *et al.*, 2014) and, therefore, calls the biological relevance of the interaction between SlKIX8 and SlTPL6 into question. To further explore this, it could be relevant to investigate the tissue-specific interactions between SlKIX and SlTPL proteins *in planta*. All in all, these data indicate that SlKIX8 and SlKIX9 are functionally redundant, but that SlKIX8 might play a predominant role in the regulation of leaf and fruit development.

### SlKIX8 and SlKIX9 are negative regulators of fruit growth

Like any other plant organ, fruit grows by means of cell division and cell expansion. After fertilization, tomato ovary growth starts with a short period of cell proliferation followed by a longer cell expansion phase, resulting in a massive expansion of the pericarp (or fruit flesh) in particular (Xiao *et al.*, 2009). Fruit ripening commences after fruit growth is finalized. Here, we report that SlKIX8 and/or SlKIX9 act as negative regulators of fruit growth, as simultaneously knocking out *SlKIX8* and *SlKIX9* by CRISRP-Cas9 genome editing results in the production of enlarged tomato fruits with increased pericarp thickness. We found that *SlKIX8* loss-of-function was sufficient to trigger an increase in tomato fruit size that was associated with an increase in pericarp thickness. A stronger effect on growth-related phenotypes by downregulation of *kix8* compared to that of *kix9* was also observed for other plant species (Gonzalez *et al.*, 2015, Nguyen *et al*., 2020). Surprisingly, the increase in tomato pericarp thickness was associated with an increased proportion of larger cells, whereas an increase in cell division was noted in the leaves of Arabidopsis *kix8 kix9* mutants. However, these Arabidopsis mutants produce larger seeds resulting from both increased cell proliferation and cell elongation (Liu *et al.*, 2020). These findings suggest that KIX8 and KIX9 might regulate multiple cellular processes, possibly through interacting with tissue-specific transcriptional regulators allowing them to target different sets of genes in function of the organ.

In line with our findings, several rosid eudicot species, in which the *KIX* or *PPD* genes were either mutated or downregulated, displayed increased seed pod size (Ge *et al.*, 2016; Kanazashi *et al.*, 2018; Li *et al.*, 2019). Ge and colleagues (Ge *et al.*, 2016) attributed the larger size of pods produced by *Medicago truncatula* plants in which an *AtPPD* ortholog was mutated to a prolonged period of cell division, similar to what they observed in developing leaves. Complete loss-of-function and severe downregulation of *PPD* genes in soybean and blackgram, respectively, led to a strong increase in seed size but was accompanied by a drastic reduction in seed number, thereby negatively impacting total yield (Kanazashi *et al.*, 2018; Naito *et al.*, 2017). Orthologs of AtSAP, an F-box protein that regulates the stability of the AtPPD-AtKIX complex, are positive regulators of seed pod and flower size in *Medicago truncatula*, of fruit and flower size in cucumber and of flower size in *Capsella rubella* (Sicard *et al.*, 2016; Yang *et al.*, 2018; Yin *et al.*, 2020).

Here, we show that tomato *kix8 kix9* loss-of-function lines produce enlarged fruits, which was among the main selection criteria for nearly all fruit crops during domestication and still is today (Pickersgill, 2007). Many of the alleles selected during domestication are not severe gain- or loss-of-function alleles, but are the result of mutations residing in *cis*-regulatory elements (CREs) that led to spatiotemporal expression changes of genes involved in crop development (Doebley *et al.*, 2006; Meyer & Purugganan, 2013; Swinnen *et al.*, 2016). These CRE alterations were likely favored over severe gain- or loss-of-function mutations, which would have been accompanied by undesirable pleiotropic effects (Swinnen *et al.*, 2016). The tomato *kix8 kix9* mutants displayed reduced axillary branching, increased flower abortion, and delayed flowering, all undesirable traits for breeding negatively impacting fruit yield. The delay in flowering time may be explained by the upregulation of *SlAP2d* in *kix8 kix9* leaves. This putative floral repressor gene is an ortholog of *AtSMZ*, which encodes a protein that counteracts the activity of CONSTANS, a promoter of flowering, by repressing the expression of multiple flowering time regulators including *FLOWERING TIME* (*FT*) in Arabidopsis leaves (Mathieu *et al.*, 2009). Engineering of CREs in the promoter regions of *SlKIX8* and *SlKIX9* to downregulate their expression specifically during fruit growth might present a more promising breeding strategy (Swinnen *et al.*, 2016; Rodríguez-Leal *et al.*, 2017, Nguyen *et al.*, 2020).

## Materials and Methods

### Ortholog identification

Tomato protein orthologs of AtKIX8 and AtKIX9 were identified through a BLASTP search in the National Center for Biotechnology Information (NCBI) GenBank protein database. Tomato protein orthologs of AtSAP, AtDFL1, AtAHL17, and AtSMZ were retrieved from the comparative genomics resource PLAZA 4.0 Dicots (http://bioinformatics.psb.ugent.be/plaza/) (Van Bel *et al.*, 2018).

### DNA Constructs

#### Yeast two- and three-hybrid constructs

For yeast two- (Y2H) and three-hybrid (Y3H) assays, the coding sequence of tomato *SlKIX8*, *SlKIX9*, *SlPPD1*, *SlPPD2*, and *SlSAP* was PCR-amplified with the primers listed in Supplemental Table 2 and recombined in a Gateway donor vector (Invitrogen). Gateway donor vectors containing the coding sequence of tomato *SlTPL1–6* were obtained from (Hao *et al.*, 2014). Subsequently, Gateway LR reactions (Invitrogen) were performed with pGAD424gate and pGBT9gate, generating bait and prey constructs, respectively. Alternatively, MultiSite Gateway LR reactions (Invitrogen) were performed with pMG426 (Nagels Durand *et al.*, 2012) to express a third protein of interest, driven by the *GDP* promoter and C-terminally fused to the SV40 NLS-3xFLAG-6xHis tag.

#### CRISPR-Cas9 constructs

To select CRISPR-Cas9 guide RNA (gRNA) target sites, CRISPR-P (http://crispr.hzau.edu.cn/CRISPR/) (Lei *et al.*, 2014) was used. We selected a gRNA target site in the first exon of *SlKIX8*, whereas for *SlKIX9*, we selected a gRNA target site in the third exon downstream of a start codon that could act as an alternative transcription start site (Supplemental Figure 2B). The CRISPR-Cas9 construct was cloned as previously described (Fauser *et al.*, 2014; Ritter *et al.*, 2017; Pauwels *et al.*, 2018). Briefly, for each gRNA target site, two complementary oligonucleotides with 4-bp overhangs (Supplemental Table 2) were annealed and inserted by a Golden Gate reaction with BpiI (Thermo Scientific) and T4 DNA ligase (Thermo Scientific) in a Gateway entry vector. As Gateway entry vectors, pMR217 (L1-R5) and pMR218 (L5-L2) (Ritter *et al.*, 2017) were used. Next, a MultiSite Gateway LR reaction (Invitrogen) was used to recombine two gRNA modules with pDe-Cas9-Km (Ritter *et al.*, 2017).

### Yeast two- and three-hybrid assays

Y2H and Y3H assays were performed as described previously (Cuéllar Pérez *et al.*, 2013). Briefly, for Y2H assays, the *Saccharomyces cerevisiae* PJ69-4A yeast strain was co-transformed with bait and prey constructs using the polyethylene glycol (PEG)/lithium acetate method. Transformants were selected on Synthetic Defined (SD) medium lacking Leu and Trp (−2) (Clontech). Three individual transformants were grown overnight in liquid SD (−2) medium and 10-fold dilutions of these cultures were dropped on SD control (−2) and selective medium additionally lacking His (−3) (Clontech). Empty vectors were used as negative controls. Yeast plates were allowed to grow for 2 days at 30°C before interaction was scored. Y3H assays were performed in the same way, but with different SD media compositions. For transformant selection and culturing in control media, SD medium lacking Leu, Trp, and Ura (−3) was used, whereas selective media additionally lacked His (−4) (Clontech).

### Plant material and growth conditions

*S. lycopersicum* wild-type and CRISPR-Cas9 mutant seeds (cultivar Micro-Tom) were sown in soil. Experiments for data in Figure 2–5, Supplemental Figure 3, Supplemental Figure 5, and Supplemental Table 1–3 were carried out in VIB-UGent (Ghent). Plants were grown under long-day photoperiods (16:8). Daytime and nighttime temperatures were 26–29°C and 18–20°C, respectively. Inflorescence and flower production, and thus fruit production, was not restricted. Experiments for data in Figure 6 and Table 1–2 were carried out in INRAE (Bordeaux). Plants were grown under long-day photoperiods (15:9). Daytime and nighttime temperatures were 25–29°C and 17–19°C, respectively. Fruit production was either unrestricted or restricted by keeping only the first two inflorescences, each carrying a maximum of 6 flowers, on the main shoot.

### Tomato plant transformation

Binary constructs were introduced in competent *Agrobacterium tumefaciens* (strain EHA105) cells using electroporation and transformed into *S. lycopersicum* (cultivar Micro-Tom) using the cotyledon transformation method as reported previously (Gonzalez *et al.*, 2007) with following modifications. Cotyledon pieces from one-week-old seedlings were incubated for 24 h in de dark at 25°C on solid Murashige and Skoog (MS) medium (pH 5.7) containing 4.4 g l^−1^ of MS supplemented with vitamins (Duchefa), 20 g l^−1^ of sucrose, 0.2 g l^−1^ of KH_2_PO_4_, 1 mg l^−1^ of thiamine, 0.2 mM of acetosyringone, 0.2 mg l^−1^ of 2,4-dichlorophenoxyacetic acid (2,4-D), and 0.1 mg l^−1^ of kinetin. Next, the cotyledon pieces were soaked in a *A. tumefaciens* (strain EHA105) bacterial suspension culture (0.05-0.10 OD) containing the binary vector for 25 min while shaking. Cotyledon pieces were dried on sterile tissue paper and placed back on the aforementioned solid MS medium for 48 h in the dark at 25°C. Cotyledon pieces were washed once with liquid MS medium (pH 5.7) containing 4.4 g l^−1^ of MS supplemented with vitamins (Duchefa), containing 20 g l^−1^ of sucrose, 0.2 g l^−1^ of KH_2_PO_4_, and 1 mg l^−1^ of thiamine and once with sterile water. Cotyledon pieces were dried on sterile tissue paper and placed on solid MS medium (pH 5.7) containing 4.4 g l^−1^ of MS supplemented with vitamins (Duchefa), containing 30 g l^−1^ of sucrose, 1 ml l^−1^ of 1000X Nitsch vitamin stock (for 100 ml: 0.005 g of biotin, 0.2 g of glycine, 10 g of myo-inositol, 0.5 g of nicotinic acid, 0.05 g of pyridoxine HCl, and 0.05 g of thiamine HCl), 0.5 g l^−1^ of folic acid, 2 mg l^−1^ of zeatin riboside, 100 mg l^−1^ of kanamycin, 25 mg l^−1^ of melatonin, and 300 mg l^−1^ of timentin and put in a 25°C controlled photoperiodic growth chamber (16:8 photoperiods). The medium was refreshed every 14 days until regenerated shoots appeared. These shoots were placed on solid MS medium (pH 5.7) containing 2.2 g l^−1^ of MS, 10 g l^− 1^ of sucrose, 1 ml l^−1^ of 1000X Nitsch vitamin stock, 0.5 g l^−1^ of folic acid, 100 mg l^−1^ of kanamycin, and 150 mg l^−1^ of timentin until their acclimatization in the greenhouse.

### Identification of CRISPR-Cas9 mutants

#### Plant genotyping

CRISPR-Cas9 mutants were identified as described previously (Swinnen *et al.*, 2020). Genomic DNA was prepared from homogenized leaf tissue using extraction buffer (pH 9.5) containing 0.1 M of tris(hydroxymethyl)aminomethane (Tris)-HCl, 0.25 M of KCl, and 0.01 M of ethylenediaminetetraacetic acid (EDTA). The mixture was incubated at 95°C for 10 min and cooled at 4°C for 5 min. After addition of 3% (w/v) BSA, the supernatant was used as a template in a standard PCR reaction using GoTaq (Promega) with Cas9-specific primers (to select primary plant transformant (T0) lines in which the T-DNA was present or plant T1 lines in which the T-DNA was absent) or with primers to amplify a gRNA target region (Supplemental Table 2). PCR amplicons containing a gRNA target site were purified using HighPrep PCR reagent (MAGBIO). After Sanger sequencing of the purified PCR amplicons with an amplification primer located approximately 200 bp from the Cas9 cleavage site, quantitative sequence trace data were decomposed using Tracking Indels by DEcomposition (TIDE) (https://www.deskgen.com/landing/tide.html#/tide) or Inference of CRISPR Editing (ICE) Analysis Tool (https://ice.synthego.com/#/).

#### Plant ploidy level analysis

Diploid CRISPR-Cas mutants (T0) were identified using flow cytometry. Leaf material (1.0 cm^2^) was chopped in 200 μl of chilled CyStain UV Precise P Nuclei Extraction Buffer (Sysmex) for 2 min using a razor blade. The suspension was filtered through a 50-μm nylon filter and 800 μl of chilled CyStain UV Precise P Staining Buffer (Sysmex) was added to the isolated nuclei. The DNA content of 5,000– 10,000 nuclei was measured using a CyFlow Space flow cytometer (Sysmex) and analyzed with FloMax software (Sysmex).

### Phenotypic analyses

#### Plant growth parameter analysis

Primary shoot, main shoot, and internode length of 4–month–old CRISPR-Cas mutant (T2) and wild-type plants was measured. Per genotype, 12 biological replicates (plants) were collected.

#### Leaf growth parameter analysis

The eighth leaf (from the top) from 2–month–old CRISPR-Cas mutant (T2) and wild-type plants was harvested for leaf growth parameter analysis. Per genotype, 31–40 biological replicates (leaves) were collected. A digital balance was used to measure the biomass/fresh weight of leaves and their terminal leaflets. Pictures of terminal leaflets were taken before (projected) and after (real) cutting the leaves to flatten them. Projected and real leaflet area were measured using ImageJ (https://imagej.nih.gov/ij/).

#### Flowering time analysis

Flowering time of CRISPR-Cas mutant (T2) and wild-type plants was quantified by counting the number of true leaves that were produced before initiation of the primary inflorescence (Soyk *et al.*, 2017). Flowering time was measured for 15–16 plants per genotype.

#### Inflorescence parameter analysis

Inflorescence and (pollinated) flower number of 3.5–4.5–month–old CRISPR-Cas mutant (T2) and wild-type plants was quantified. Per genotype, 12 biological replicates (plants) were collected.

#### Ovary growth parameter analysis

For ovary diameter and height measurements, ovaries from CRISPR-Cas mutant (T2) and wild-type plants were harvested at anthesis. Ovary diameter was determined by averaging the maximum and minimum diameter of the equatorial axis. Per genotype, 3 biological replicates (ovaries) were collected.

#### Fruit growth parameter analysis

For fruit biomass, pericarp thickness, and yield measurements, fruits at distinct developmental and ripening stages (5 days post anthesis–red ripe) were harvested from CRISPR-Cas mutant (T2) and wild-type plants. Pericarp thickness was measured by taking scans of equatorial fruit sections and using Tomato Analyzer (version 4.0) (Brewer *et al.*, 2006). Per genotype and developmental stage, 12 biological replicates (fruits from 12 individual plants; fruit production unrestricted; VIB-UGent), 7–9 biological replicates (fruits from 12 individual plants; fruit production unrestricted; INRAE) or 3 biological replicates (fruits from 3 batches of 3 plants; fruit production restricted; INRAE) were collected.

For cell layer number quantification and cell area measurements, fruits were harvested at 30 days post anthesis and pericarp was fixed in a solution of FAA (18v EtOH 70%, 1v acetic acid, and 1v formaldehyde). Pericarp sections with a thickness of 100 μm were made with a Microtom and imaged. The number of cell layers and individual cell area were quantified using ImageJ (https://imagej.nih.gov/ij/). Per genotype, 2–4 biological replicates (fruit production restricted; INRAE) were collected.

#### Seed parameter analysis

For seed number and size analyses, seeds were harvested from red ripe fruits produced by CRISPR-Cas mutant (T2) and wild-type plants. Seed area was measured using ImageJ (https://imagej.nih.gov/ij/). Per genotype, 12 biological replicates were collected, constituting seeds from red ripe fruits from 12 individual plants.

#### Statistical analysis

For all phenotypic analyses, statistical significance was determined by ANOVA followed by Tukey post-hoc analysis (P < 0.05).

### Gene expression analysis by quantitative real-time PCR

The terminal leaflet of the second leaf (from the top) from 3-week-old CRISPR-Cas mutant (T2) and wild-type plants was harvested by flash freezing in liquid nitrogen and ground using the Mixer Mill 300 (Retch). Per genotype, five biological replicates, each consisting of a single terminal leaflet were collected. Messenger RNA was extracted from homogenized tissue as described in (Townsley *et al.*, 2015) with following modifications. Tissue was lysed using 800 μl of lysate binding buffer (LBB) containing 100 mM of Tris-HCl (pH 7.5), 500 mM of LiCl, 10 mM of EDTA (pH 8.0), 1% of sodium dodecyl sulfate (SDS), 5 mM of dithiothreitol (DTT), 15 μl ml^−1^ of Antifoam A, and 5 μl ml^−1^ of 2-mercaptoethanol, and the mixture was incubated for 10 min at room temperature. Messenger RNA was separated from 200 μl of lysate using 1 μl of 12.5 μM of 5’ biotinylated polyT oligonucleotide (5’-biotin- ACAGGACATTCGTCGCTTCCTTTTTTTTTTTTTTTTTTTT-3’) and the mixture was incubated for 10 min. Next, captured messenger RNA was isolated from the lysate by adding 20 μl of LBB-washed streptavidin-coated magnetic beads (New England Biolabs) and incubated for 10 min at room temperature. Samples were placed on a MagWell Magnetic Separator 96 (EdgeBio) and washed with 200 μl of washing buffer A (10 mM of Tris-HCl (pH 7.5), 150 mM of LiCl, 1 mM of EDTA (pH 8.0), 0.1% of SDS), washing buffer B (10 mM of Tris-HCl (pH 7.5), 150 mM of LiCl, 1 mM of EDTA (pH 8.0)), and low-salt buffer (20 mM of Tris-HCl (pH 7.5), 150 mM of NaCl, 1 mM of EDTA (pH 8.0)), all pre-chilled on ice. Elution of messenger RNA was done by adding 20 μl of 10 mM of Tris-HCl (pH 8.0) with 1 mM of 2-mercaptoethanol followed by incubation of the mixture at 80°C for 2 min.

First-strand complementary DNA was synthesized from 20 μl of messenger RNA eluate by qScript cDNA Synthesis Kit (Quantabio). Quantitative real-time PCR (qPCR) reactions were performed with a LightCycler 480 System (Roche) using Fast SYBR Green Master Mix (Applied Biosystems) and primers (Supplemental Table 2) designed by QuantPrime (https://www.quantprime.de/) (Arvidsson *et al.*, 2008). Gene expression levels were quantified relative to *CLATHRIN ADAPTOR COMPLEXES MEDIUM SUBUNIT* (*SlCAC*) and *TAP42-INTERACTING PROTEIN* (*SlTIP41*) using the 2^−ΔΔCt^ method (Livak & Schmittgen, 2001). Statistical significance was determined by ANOVA followed by Tukey post-hoc analysis (P < 0.05).

## Accession numbers

Sequence data from this article can be found in the EMBL/GenBank/Solgenomics data libraries under the following accession numbers: *SlKIX8* (Solyc07g008100), *SlKIX9* (Solyc08g059700), *SlPPD1* (Solyc06g084120), *SlPPD2* (Solyc09g065630), *SlSAP* (Solyc05g041220), *SlDFL1* (Solyc07g063850), *SlAHL17* (Solyc04g076220), *SlAP2d* (Solyc11g072600), *SlCAC* (Solyc08g006960), and *SlTIP41* (Solyc10g049850).

## Acknowledgements

This work was supported by the Research Foundation Flanders (FWO) through the projects G005312N, G004515N, and 3G038719 and a postdoctoral fellowship to L.P. We thank Annick Bleys for help with preparing the manuscript and Mohamed Zouine for sharing plasmids containing the coding sequence of *SlTPL1–6* with us.

## Supplemental Materials

**Supplemental Figure 1.** A conserved repressor complex regulates leaf growth in distinct eudicot species.

**Supplemental Figure 2.** Splice variants of *SlKIX8*, *SlKIX9*, *SlPPD1*, and *SlPPD2*.

**Supplemental Figure 3.** Regenerated tomato *kix8 kix9* plants display a rippled, dome-shaped leaf phenotype.

**Supplemental Figure 4.** CRISPR-Cas9 mutations in double *kix8 kix9* (T1), single *kix8*, and single *kix9* tomato knockout lines.

**Supplemental Figure 5.** Tomato *kix8 kix9* plants display a delay in flowering time.

**Supplemental Table 1.** Tomato *kix8 kix9* plants display a reduction in plant height.

**Supplemental Table 2.** Normalized expression of *SlKIX8*, *SlKIX9*, *SlPPD1*, *SlPPD2*, *SlDFL1*, *SlAHL17*, and *SlAP2d* in different tomato organs and developmental stages (cultivar Micro-Tom) used to generate heat maps in Figure 4, A and B.

**Supplemental Table 3.** Tomato *kix8 kix9* plants display a reduction in in axillary shoot formation.

**Supplemental Table 4.** Oligonucleotides used in this study.

## Supplemental Materials

**Figure S1.**
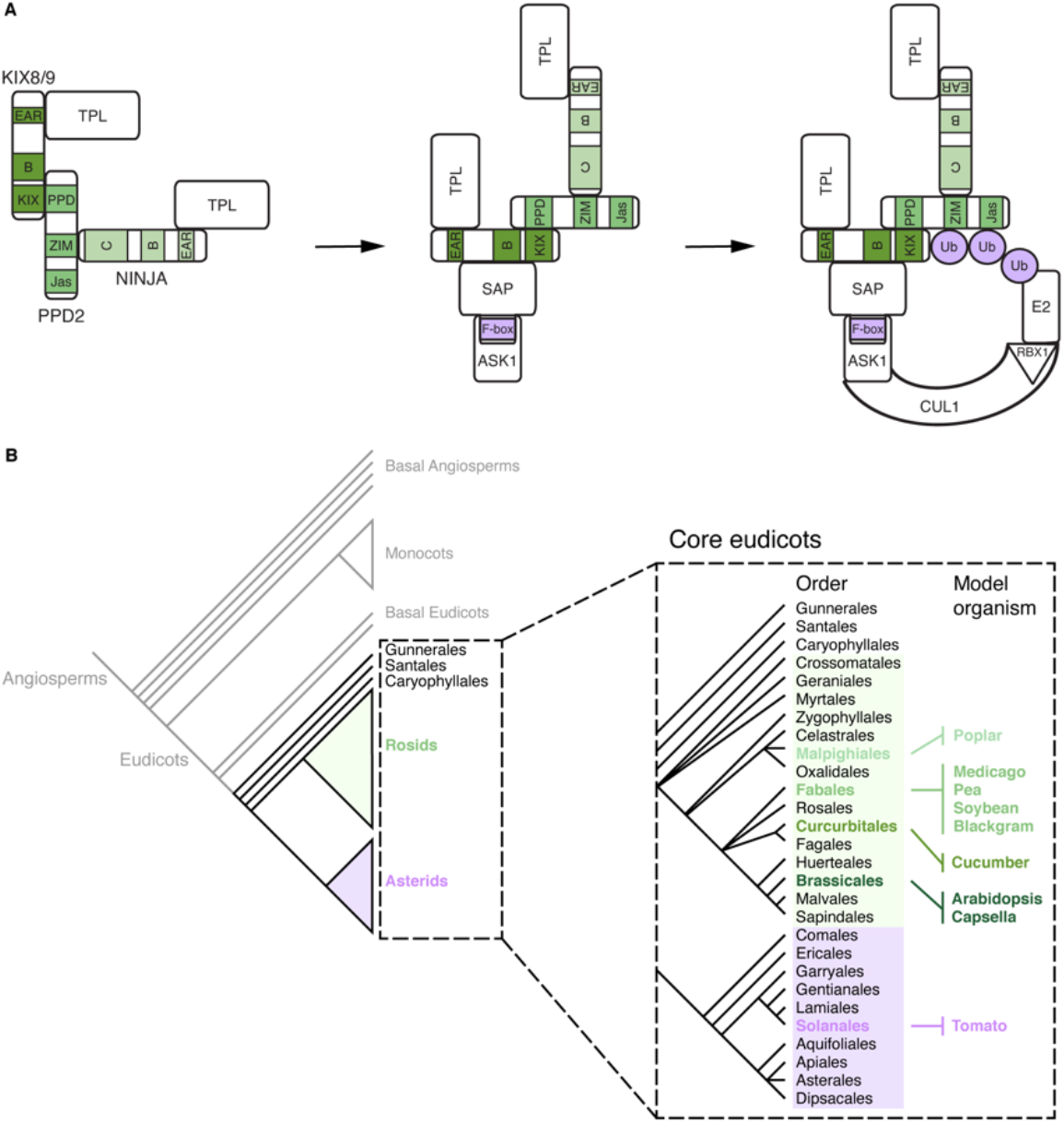
A conserved repressor complex regulates leaf growth in distinct eudicot species. (A) The AtPPD2-AtKIX8/AtKIX9 transcriptional repressor complex in *Arabidopsis thaliana*. AtPPD2 interacts with AtKIX8/AtKIX9 and AtNINJA to recruit AtTPL. Interaction of repressor complex members with the E3 ubiquitin ligase AtSCF^SAP^ (comprising the F-box protein AtSAP, AtASK1, AtCUL1, and AtRBX1) leads to the proteasomal degradation of AtKIX8/AtKIX9 and AtPPD2. (B) Model organisms in which KIX, PPD and/or SAP proteins were shown to mediate leaf growth belong to different orders within the rosids, which together with the asterids, make up most of the core eudicot species. Tomato is an asterid model species in which the potential role of these proteins in regulating leaf growth has not been investigated yet. Abbreviations: ASK1, Arabidopsis SKP1; CUL1, CULLIN 1; RBX1, RING-BOX 1; SKP1, S-PHASE KINASE-ASSOCIATED PROTEIN 1; Ub, ubiquitin.

**Figure S2.**
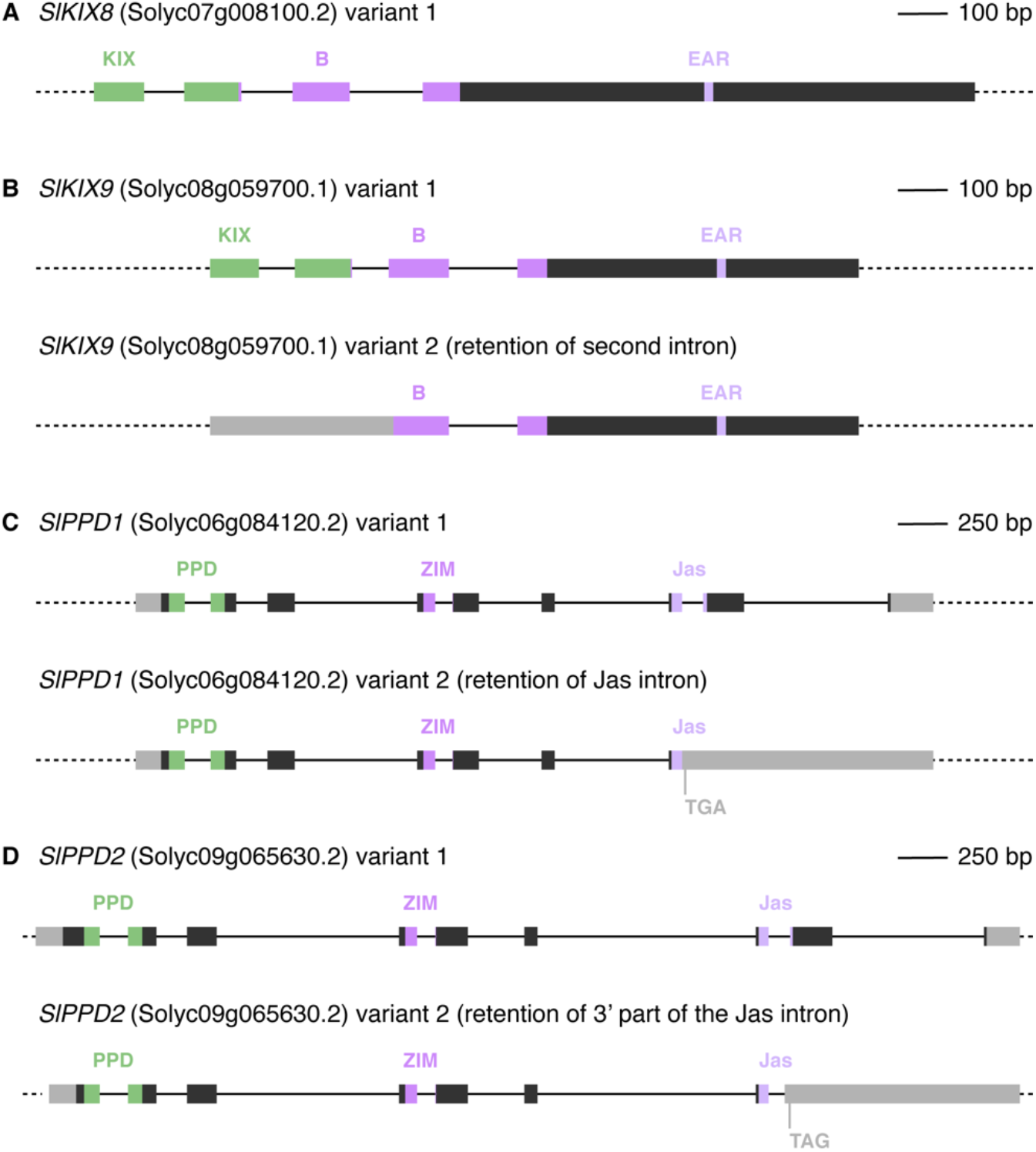
Splice variants of *SlKIX8*, *SlKIX9*, *SlPPD1*, and *SlPPD2*. Dark grey boxes represent exons, solid lines represent introns and light grey boxes represent UTRs. Green and purple boxes represent encoded protein domains. No alternative splicing was observed for *SlKIX8* (A). Retention of the second *SlKIX9* intron (B) could lead to the use of a downstream start codon, excluding the sequence that encodes the N-terminal KIX domain. The splice variants of *SlPPD1* (C) and *SlPPD2* (D) display retention of the Jas intron and part of the Jas intron, respectively, which is located between the two exons that encode the Jas domain. These alternative splicing events generate premature stop codons. Abbreviations: UTR, untranslated region.

**Figure S3.**
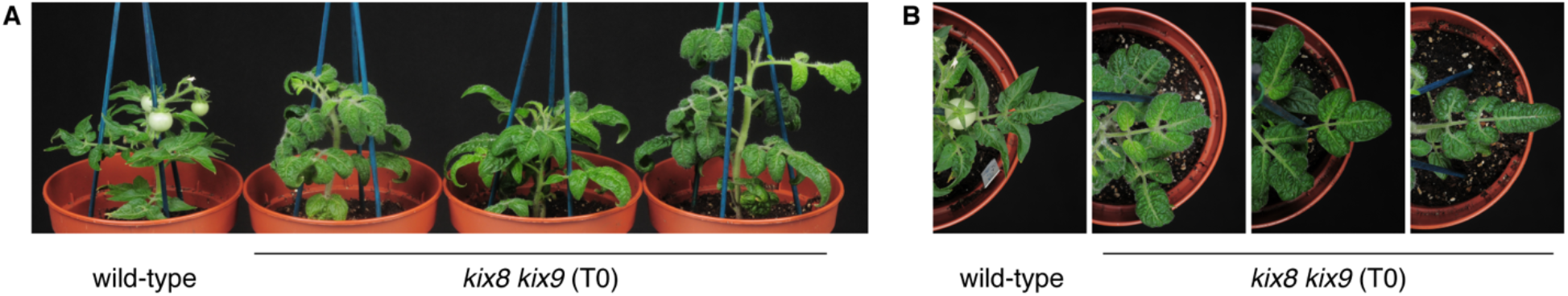
Regenerated tomato *kix8 kix9* plants display a rippled, dome-shaped leaf phenotype. (A−B) Wild-type and regenerated *kix8 kix9* plants were photographed from the front (A) and the top (B). Primary transformants transferred from rooting medium were grown in soil for 10 weeks under 16:8 photoperiods with daytime and nighttime temperatures of 26–29°C and 18–20°C, respectively.

**Figure S4.**
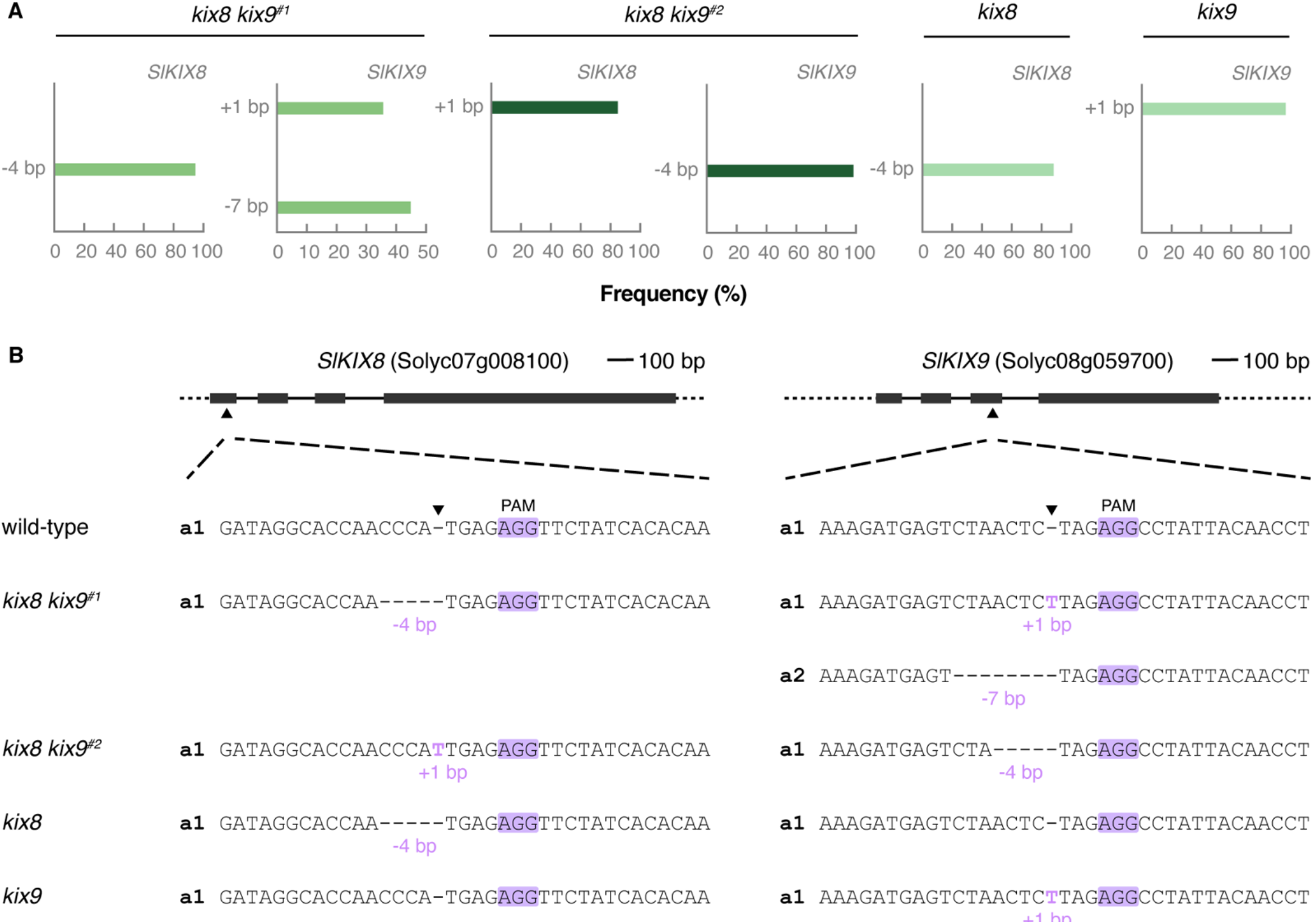
CRISPR-Cas9 mutations in double *kix8 kix9* (T1) and single *kix8*, and single *kix9* tomato knockout lines. (A) ICE analysis of genomic sites targeted by the guide RNAs. Targeted genomic regions were PCR amplified and sequenced by Sanger sequencing. Based on the sequence chromatograms, ICE analysis visualized the indel spectrum and calculated the frequency of each indel. (B) Schematic representation of *SlKIX8* and *SlKIX9* with location of the CRISPR-Cas9 cleavage sites. Dark grey boxes represent exons and solid lines represent introns. Cas9 cleavage sites for guide RNAs are indicated with arrowheads. Allele sequences are shown for two independent *kix8 kix9* lines, one *kix8* line, and one *kix9* line. Abbreviations: a, allele; PAM, protospacer adjacent motif.

**Figure S5.**
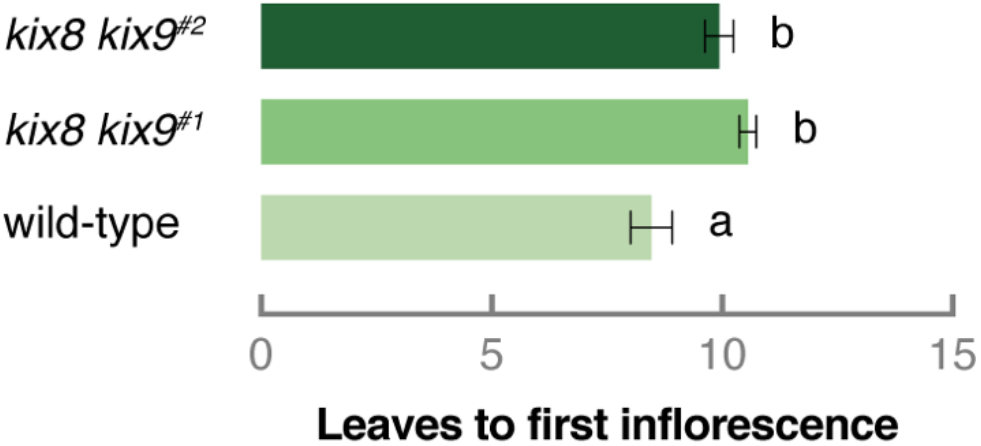
Tomato *kix8 kix9* plants display a delay in flowering time. Plants were grown in soil under 16:8 photoperiods with daytime and nighttime temperatures of 26–29°C and 18–20°C, respectively. Bars represent mean number of leaves produced before initiation of the first inflorescence. Error bars denote standard error (n = 15–16). Statistical significance was determined by ANOVA followed by Tukey post-hoc analysis (P < 0.05; indicated by different letters).

**Table S1.**
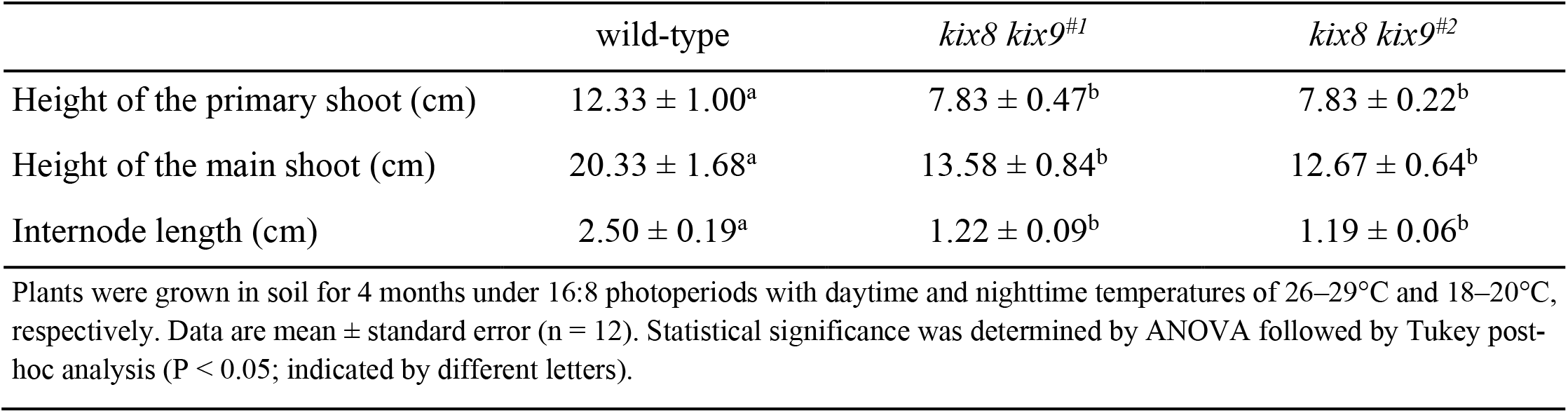
Tomato *kix8 kix9* plants display a reduction in plant height.

**Table S2.**
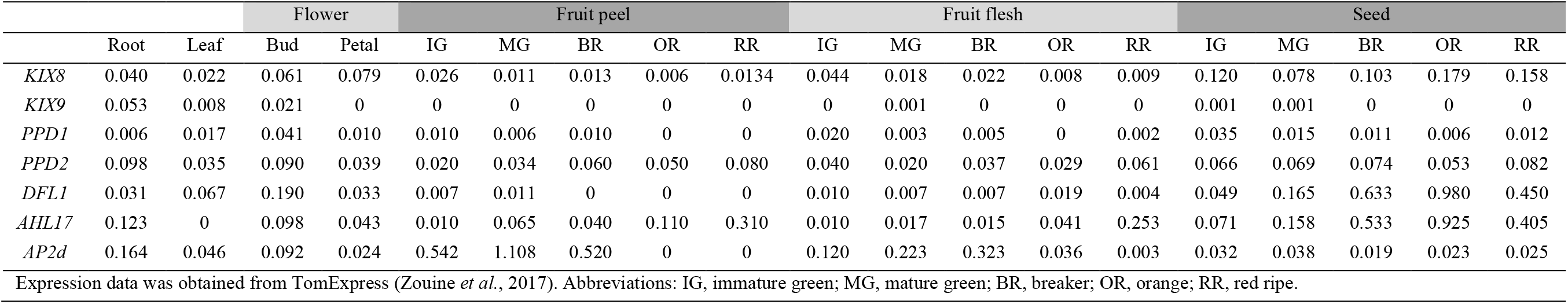
Normalized expression of *SlKIX8*, *SlKIX9*, *SlPPD1*, *SlPPD2*, *SlDFL1*, *SlAHL17*, and *SlAP2d* in different tomato organs and developmental stages (cultivar Micro-Tom) used to generate heat maps in Figure 4, A and B.

**Table S3.**
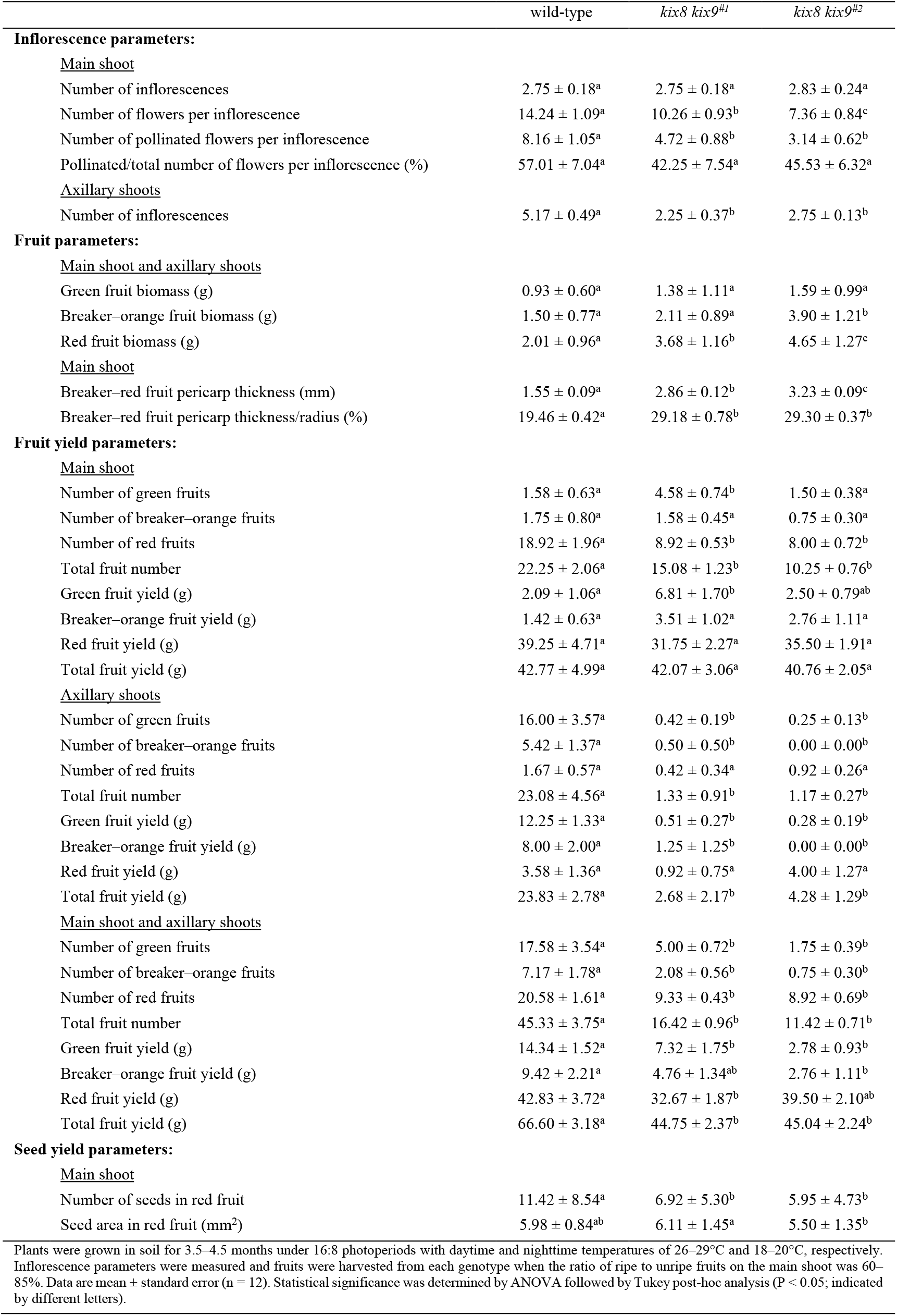
Tomato *kix8 kix9* plants display a reduction in axillary shoot formation

**Table S4.**
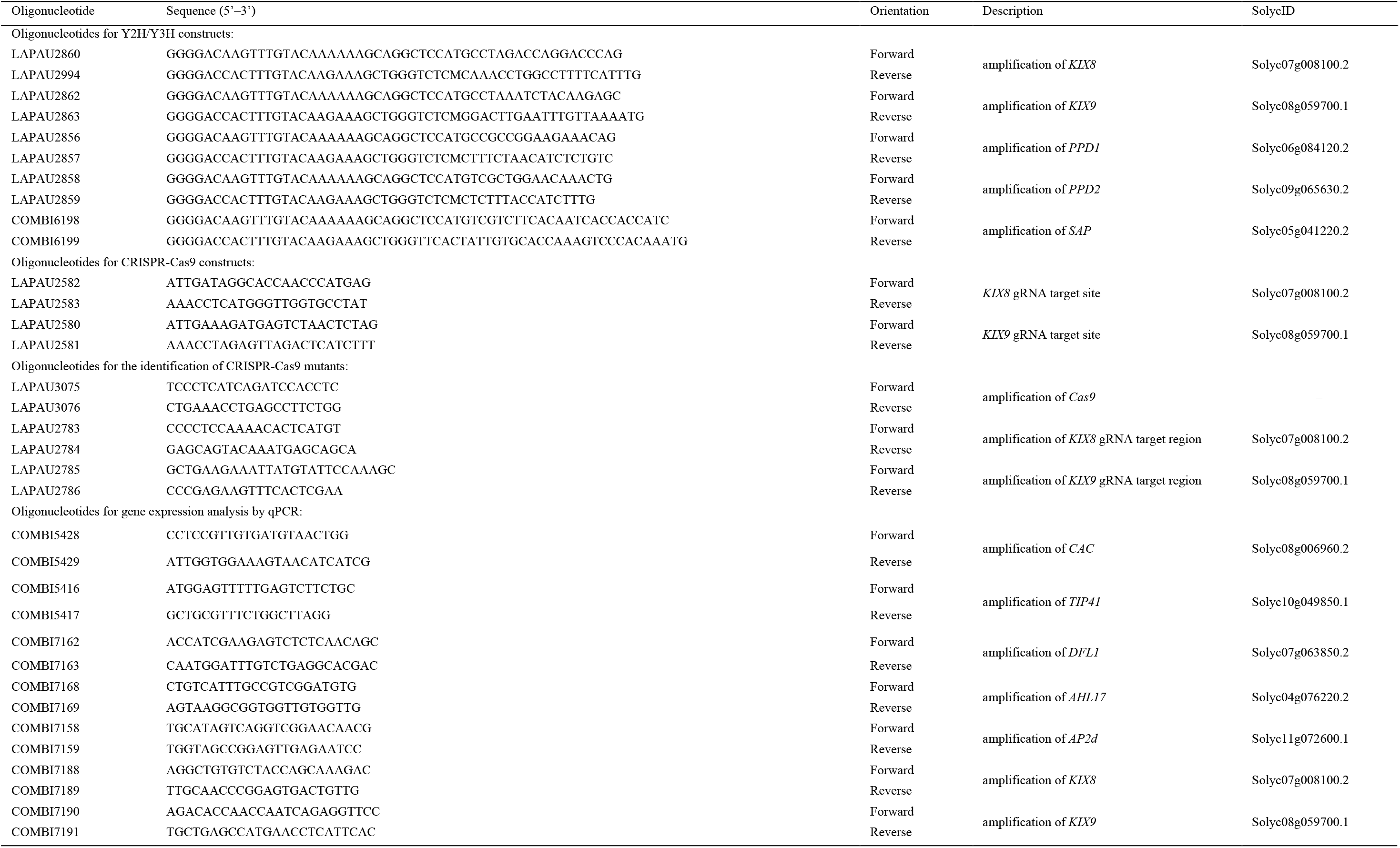
Oligonucleotides used in this study.

## References

Arvidsson S, Kwasniewski M, Riaño-Pachón DM, Mueller-Roeber B. 2008. QuantPrime - a flexible tool for reliable high-throughput primer design for quantitative PCR. BMC Bioinformatics 9: 465.

Baekelandt A, Pauwels L, Wang ZB, Li N, De Milde L, Natran A, Vermeersch M, Li Y, Goossens A, Inzé D, et al. 2018. Arabidopsis leaf flatness is regulated by PPD2 and NINJA through repression of *CYCLIN D3* genes. Plant Physiology 178: 217–232.

Bai Y, Meng Y, Huang D, Qi Y, Chen M. 2011. Origin and evolutionary analysis of the plant-specific TIFY transcription factor family. Genomics 98: 128–136.

Brewer MT, Lang L, Fujimura K, Dujmovic N, Gray S, van der Knaap E. 2006. Development of a controlled vocabulary and software application to analyze fruit shape variation in tomato and other plant species. Plant Physiology 141: 15–25.

Causier B, Ashworth M, Guo W, Davies B. 2012. The TOPLESS interactome: a framework for gene repression in Arabidopsis. Plant Physiology 158: 423–438.

Chini A, Ben-Romdhane W, Hassairi A, Aboul-Soud MAM. 2017. Identification of TIFY/JAZ family genes in *Solanum lycopersicum* and their regulation in response to abiotic stresses. PLoS ONE 12: e0177381.

Chini A, Fonseca S, Chico JM, Fernández-Calvo P, Solano R. 2009. The ZIM domain mediates homo- and heteromeric interactions between Arabidopsis JAZ proteins. Plant Journal 59: 77–87.

Chini A, Fonseca S, Fernández G, Adie B, Chico JM, Lorenzo O, García-Casado G, López-Vidriero I, Lozano FM, Ponce MR, et al. 2007. The JAZ family of repressors is the missing link in jasmonate signalling. Nature 448: 666–671.

Chung HS, Howe GA. 2009. A critical role for the TIFY motif in repression of jasmonate signaling by a stabilized splice variant of the JASMONATE ZIM-domain protein JAZ10 in *Arabidopsis*. Plant Cell 21: 131–145.

Cuéllar Pérez A, Pauwels L, De Clercq R, Goossens A. 2013. Yeast two-hybrid analysis of jasmonate signaling proteins. Methods in Molecular Biology 1011: 173–185.

Doebley JF, Gaut BS, Smith BD. 2006. The molecular genetics of crop domestication. Cell 127: 1309–1321.

Fauser F, Schiml S, Puchta H. 2014. Both CRISPR/Cas-based nucleases and nickases can be used efficiently for genome engineering in *Arabidopsis thaliana*. Plant Journal 79: 348–359.

Ge L, Yu J, Wang H, Luth D, Bai G, Wang K, Chen R. 2016. Increasing seed size and quality by manipulating *BIG SEEDS1* in legume species. Proceedings of the National Academy of Sciences of the United States of America 113: 12414–12419.

Gonzalez N, Gévaudant F, Hernould M, Chevalier C, Mouras A. 2007. The cell cycle-associated protein kinase WEE1 regulates cell size in relation to endoreduplication in developing tomato fruit. Plant Journal 51: 642–655.

Gonzalez N, Pauwels L, Baekelandt A, De Milde L, Van Leene J, Besbrugge N, Heyndrickx KS, Cuéllar Pérez A, Nagels Durand A, De Clercq R, et al. 2015. A repressor protein complex regulates leaf growth in Arabidopsis. Plant Cell 27: 2273–2287.

Gonzalez N, Vanhaeren H, Inzé D. 2012. Leaf size control: complex coordination of cell division and expansion. Trends in Plant Science 17: 332–340.

Hao Y, Wang X, Li X, Bassa C, Mila I, Audran C, Maza E, Li Z, Bouzayen M, van der Rest B, et al. 2014. Genome-wide identification, phylogenetic analysis, expression profiling, and protein–protein interaction properties of *TOPLESS* gene family members in tomato. Journal of Experimental Botany 65: 1013–1023.

Hepworth J, Lenhard M. 2014. Regulation of plant lateral-organ growth by modulating cell number and size. Current Opinion in Plant Biology 17: 36–42.

Kagale S, Links MG, Rozwadowski K. 2010. Genome-wide analysis of ethylene-responsive element binding factor-associated amphiphilic repression motif-containing transcriptional regulators in Arabidopsis. Plant Physiology 152: 1109–1134.

Kajala K, Shaar-Moshe L, Mason GA, Gouran M, Rodriguez-Medina J, Kawa D, Pauluzzi G, Reynoso M, Canto-Pastor A, Lau V, et al. 2020. Innovation, conservation and repurposing of gene function in plant root cell type development. bioRxiv: 2020.2004.2009.017285.

Kalve S, De Vos D, Beemster GTS. 2014. Leaf development: a cellular perspective. Frontiers in Plant Science 5: 362.

Kanazashi Y, Hirose A, Takahashi I, Mikami M, Endo M, Hirose S, Toki S, Kaga A, Naito K, Ishimoto M, et al. 2018. Simultaneous site-directed mutagenesis of duplicated loci in soybean using a single guide RNA. Plant Cell Reports. doi: 10.1007/s00299-018-2251-3

Kumar V, Waseem M, Dwivedi N, Maji S, Kumar A, Thakur JK. 2018. KIX domain of AtMed15a, a Mediator subunit of Arabidopsis, is required for its interaction with different proteins. Plant Signaling & Behavior 13: e1428514.

Lei Y, Lu L, Liu H-Y, Li S, Xing F, Chen L-L. 2014. CRISPR-P: a web tool for synthetic single-guide RNA design of CRISPR-system in plants. Molecular Plant 7: 1494–1496.

Li N, Liu Z, Wang Z, Ru L, Gonzalez N, Baekelandt A, Pauwels L, Goossens A, Xu R, Zu Z, et al. 2018. STERILE APETALA modulates the stability of a repressor protein complex to control organ size in *Arabidopsis thaliana*. PLoS Genetics 14: e1007218.

Li S, Yamada M, Hang X, Ohler U, Benfey PN. 2016. High-resolution expression map of the *Arabidopsis* root reveals alternative splicing and lincRNA regulation. Developmental Cell 39: 508–522.

Li X, Liu W, Zhuang L, Zhu Y, Wang F, Chen T, Yang J, Ambrose M, Hu Z, Weller JL, et al. 2019. BIGGER ORGANS and ELEPHANT EAR-LIKE LEAF1 control organ size and floral organ internal asymmetry in pea. Journal of Experimental Botany 70: 179–191.

Liu Z, Na L, Zhang Y, Li Y. 2020. Transcriptional repression of *GIF1* by the KIX-PPD-MYC repressor complex controls seed size in Arabidopsis. Nature Communications 11: 1846.

Liu T, Ohashi-Ito K, Bergmann DC. 2009. Orthologs of *Arabidopsis thaliana* stomatal bHLH genes and regulation of stomatal development in grasses. Development 136: 2265–2276.

Livak KJ, Schmittgen TD. 2001. Analysis of relative gene expression data using real-time quantitative PCR and the 2^−ΔΔCT^ method. Methods 25: 402–408.

Mathieu J, Yant LJ, Murdter F, Küttner F, Schmid M. 2009. Repression of flowering by the miR172 target SMZ. PLoS Biology 7: e1000148.

Meyer RS, Purugganan MD. 2013. Evolution of crop species: genetics of domestication and diversification. Nature Reviews Genetics 14: 840–852.

Nagels Durand A, Moses T, De Clercq R, Goossens A, Pauwels L. 2012. A MultiSite Gateway™ vector set for the functional analysis of genes in the model *Saccharomyces cerevisiae*. BMC Molecular Biology 13: 30.

Naito K, Takahashi Y, Chaitieng B, Hirano K, Kaga A, Takagi K, Ogiso-Tanaka E, Thavarasook C, Ishimoto M, Tomooka N. 2017. Multiple organ gigantism caused by mutation in *VmPPD* gene in blackgram (*Vigna mungo*). Breeding Science 67: 151–158.

Nelissen H, Moloney M, Inzé D. 2014. Translational research: from pot to plot. Plant Biotechnology Journal 12: 277–285.

Nguyen CX, Paddock KJ, Zhang Z, Stacey MG. 2021. GmKIX8-1 regulates organ size in soybean and is the causative gene for the major seed weight QTL *qSw17-1*. New Phytologist 2: 920–934.

Pauwels L, Barbero GF, Geerinck J, Tilleman S, Grunewald W, Cuéllar Pérez A, Chico JM, Vanden Bossche R, Sewell J, Gil E, et al. 2010. NINJA connects the co-repressor TOPLESS to jasmonate signalling. Nature 464: 788–791.

Pauwels L, De Clercq R, Goossens J, Iñigo S, Williams C, Ron M, Britt A, Goossens A. 2018. A dual sgRNA approach for functional genomics in *Arabidopsis thaliana*. G3: Genes, Genomes, Genetics 8: 2603–2615.

Pickersgill B. 2007. Domestication of plants in the Americas: insights from mendelian and molecular genetics. Annals of Botany 100: 925–940.

Ritter A, Iñigo S, Fernández-Calvo P, Heyndrickx KS, Dhondt S, Shi H, De Milde L, Vanden Bossche R, De Clercq R, Eeckhout D, et al. 2017. The transcriptional repressor complex FRS7-FRS12 regulates flowering time and growth in *Arabidopsis*. Nature Communications 8: 15235.

Rodríguez-Leal D, Lemmon ZH, Man J, Bartlett ME, Lippman ZB. 2017. Engineering quantitative trait variation for crop improvement by genome editing. Cell 171: 470–480.e478.

Schneider M, Gonzalez N, Pauwels L, Inzé D, Baekelandt A. 2021. The PEAPOD Pathway and Its Potential To Improve Crop Yield. Trends Plant Sci 26: 220–236

Sicard A, Kappel C, Lee YW, Woźniak, N. J., Marona C, Stinchcombe JR, Wright SI, Lenhard M. 2016. Standing genetic variation in a tissue-specific enhancer underlies selfing-syndrome evolution in *Capsella*. Proceedings of the National Academy of Sciences of the United States of America 113: 13911–13916.

Soyk S, Müller NA, Park SJ, Schmalenbach I, Jiang K, Hayama R, Zhang L, Van Eck J, Jiménez-Gómez JM, Lippman ZB. 2017. Variation in the flowering gene *SELF PRUNING 5G* promotes day-neutrality and early yield in tomato. Nature Genetics 49: 162–168.

Swinnen G, Goossens A, Pauwels L. 2016. Lessons from domestication: targeting *cis*-regulatory elements for crop improvement. Trends in Plant Science 21: 506–515.

Swinnen G, Jacobs T, Pauwels L, Goossens A. 2020. CRISPR-Cas-mediated gene knockout in tomato. Methods in Molecular Biology 2083: 321–341.

Thakur JK, Agarwal P, Parida S, Bajaj D, Pasrija R. 2013. Sequence and expression analyses of KIX domain proteins suggest their importance in seed development and determination of seed size in rice, and genome stability in Arabidopsis. Molecular Genetics and Genomics 288: 329–346.

Thakur JK, Yadav A, Yadav G. 2014. Molecular recognition by the KIX domain and its role in gene regulation. Nucleic Acids Research 42: 2112–2125.

Thines B, Katsir L, Melotto M, Niu Y, Mandaokar A, Liu G, Nomura K, He SY, Howe GA, Browse J. 2007. JAZ repressor proteins are targets of the SCF^COI1^ complex during jasmonate signalling. Nature 448: 661–665.

Townsley BT, Covington MF, Ichihashi Y, Zumstein K, Sinha NR. 2015. BrAD-seq: Breath Adapter Directional sequencing: a streamlined, ultra-simple and fast library preparation protocol for strand specific mRNA library construction. Frontiers in Plant Science 6: 366.

Van Bel M, Diels T, Vancaester E, Kreft L, Botzki A, Van de Peer Y, Coppens F, Vandepoele K. 2018. PLAZA 4.0: an integrative resource for functional, evolutionary and comparative plant genomics. Nucleic Acids Research 46: D1190–D1196.

Vanholme B, Grunewald W, Bateman A, Kohchi T, Gheysen G. 2007. The tify family previously known as ZIM. Trends in Plant Science 12: 239–244.

Vatén A, Bergmann DC. 2012. Mechanisms of stomatal development: an evolutionary view. EvoDevo 3: 11.

Vercruysse J, Baekelandt A, Gonzalez N, Inzé D. 2020. Molecular networks regulating the cell division during leaf growth in Arabidopsis. Journal of Experimental Botany. doi: 10.1093/jxb/erz522

Wang Z, Li N, Jiang S, Gonzalez N, Huang X, Wang Y, Inzé D, Li Y. 2016. SCF^SAP^ controls organ size by targeting PPD proteins for degradation in *Arabidopsis thaliana*. Nature Communications 7: 11192.

White DWR. 2006. *PEAPOD* regulates lamina size and curvature in *Arabidopsis*. Proceedings of the National Academy of Sciences of the United States of America 103: 13238–13243.

Wikström N, Savolainen V, Chase MW. 2001. Evolution of the angiosperms: calibrating the family tree. Proceedings of the Royal Society of London Series B-Biological Sciences 268: 2211–2220.

Xiao H, Radovich C, Welty N, Hsu J, Li D, Meulia T, van der Knaap E. 2009. Integration of tomato reproductive developmental landmarks and expression profiles, and the effect of *SUN* on fruit shape. BMC Plant Biology 9: 49.

Yang L, Liu H, Zhao J, Pan Y, Cheng S, Lietzow CD, Wen C, Zhang X, Weng Y. 2018. *LITTLELEAF* (*LL*) encodes a WD40 repeat domain-containing protein associated with organ size variation in cucumber. Plant Journal 95: 834–847.

Yin P, Ma Q, Wang H, Feng D, Wang X, Pei Y, Wen J, Tadege M, Niu L, Lin H. 2020. SMALL LEAF AND BUSHY1 controls organ size and lateral branching by modulating the stability of BIG SEEDS1 in *Medicago truncatula*. New Phytologist. doi: 10.1111/nph.16449

Yordanov YS, Ma C, Yordanova E, Meilan R, Strauss SH, Busov VB. 2017. *BIG LEAF* is a regulator of organ size and adventitious root formation in poplar. PLoS ONE 12: e0180527.

Zouine M, Maza E, Djari A, Lauvernier M, Frasse P, Smouni A, Pirrello J, Bouzayen M. 2017. TomExpress, a unified tomato RNA-Seq platform for visualization of expression data, clustering and correlation networks. Plant Journal 92: 727–735.

