## Supplemental Materials for "SlKIX8 and SlKIX9 are negative regulators of leaf and fruit growth in tomato"

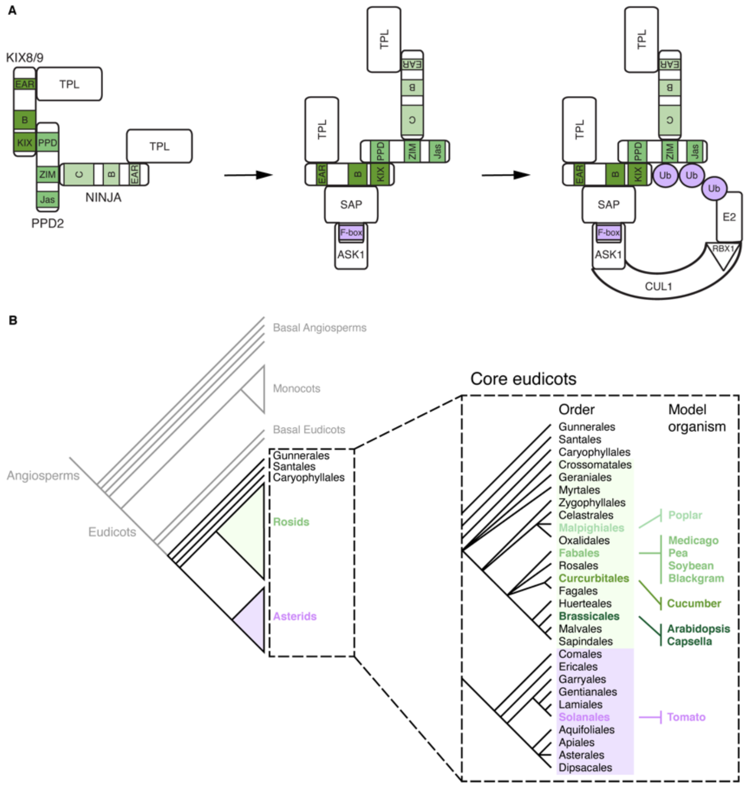


**Figure S1.** A conserved repressor complex regulates leaf growth in distinct eudicot species. (A) The AtPPD2-AtKIX8/AtKIX9 transcriptional repressor complex in *Arabidopsis thaliana*. AtPPD2 interacts with AtKIX8/AtKIX9 and AtNINJA to recruit AtTPL. Interaction of repressor complex members with the E3 ubiquitin ligase $AtSCF^{SAP}$ (comprising the F-box protein AtSAP, AtASK1, AtCUL1, and AtRBX1) leads to the proteasomal degradation of AtKIX8/AtKIX9 and AtPPD2. (B) Model organisms in which KIX, PPD and/or SAP proteins were shown to mediate leaf growth belong to different orders within the rosids, which together with the asterids, make up most of the core eudicot species. Tomato is an asterid model species in which the potential role of these proteins in regulating leaf growth has not been investigated yet. Abbreviations: ASK1, Arabidopsis SKP1; CUL1, CULLIN 1; RBX1, RING-BOX 1; SKP1, S-PHASE KINASE-ASSOCIATED PROTEIN 1; Ub, ubiquitin.

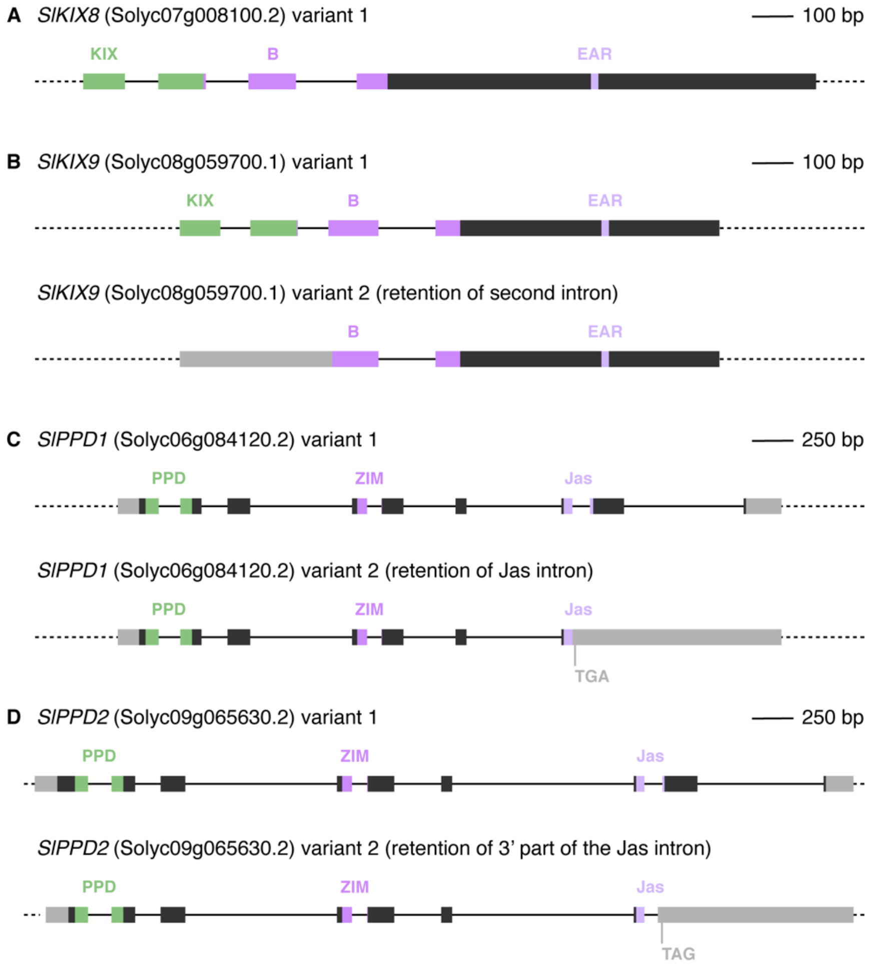


**Figure S2.** Splice variants of *SlKIX8*, *SlKIX9*, *SlPPD1*, and *SlPPD2*. Dark grey boxes represent exons, solid lines represent introns and light grey boxes represent UTRs. Green and purple boxes represent encoded protein domains. No alternative splicing was observed for *SlKIX8* (A). Retention of the second *SlKIX9* intron (B) could lead to the use of a downstream start codon, excluding the sequence that encodes the N-terminal KIX domain. The splice variants of *SlPPD1* (C) and *SlPPD2* (D) display retention of the Jas intron and part of the Jas intron, respectively, which is located between the two exons that encode the Jas domain. These alternative splicing events generate premature stop codons. Abbreviations: UTR, untranslated region.

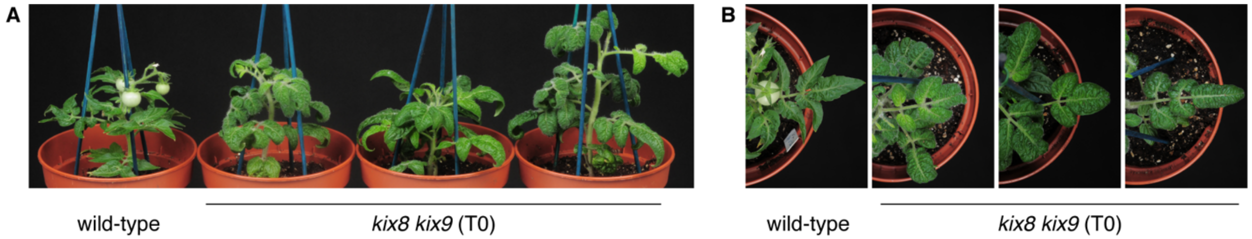


**Figure S3.** Regenerated tomato *kix8 kix9* plants display a rippled, dome-shaped leaf phenotype. (A−B) Wild-type and regenerated *kix8 kix9* plants were photographed from the front (A) and the top (B). Primary transformants transferred from rooting medium were grown in soil for 10 weeks under 16:8 photoperiods with daytime and nighttime temperatures of 26–29°C and 18–20°C, respectively.

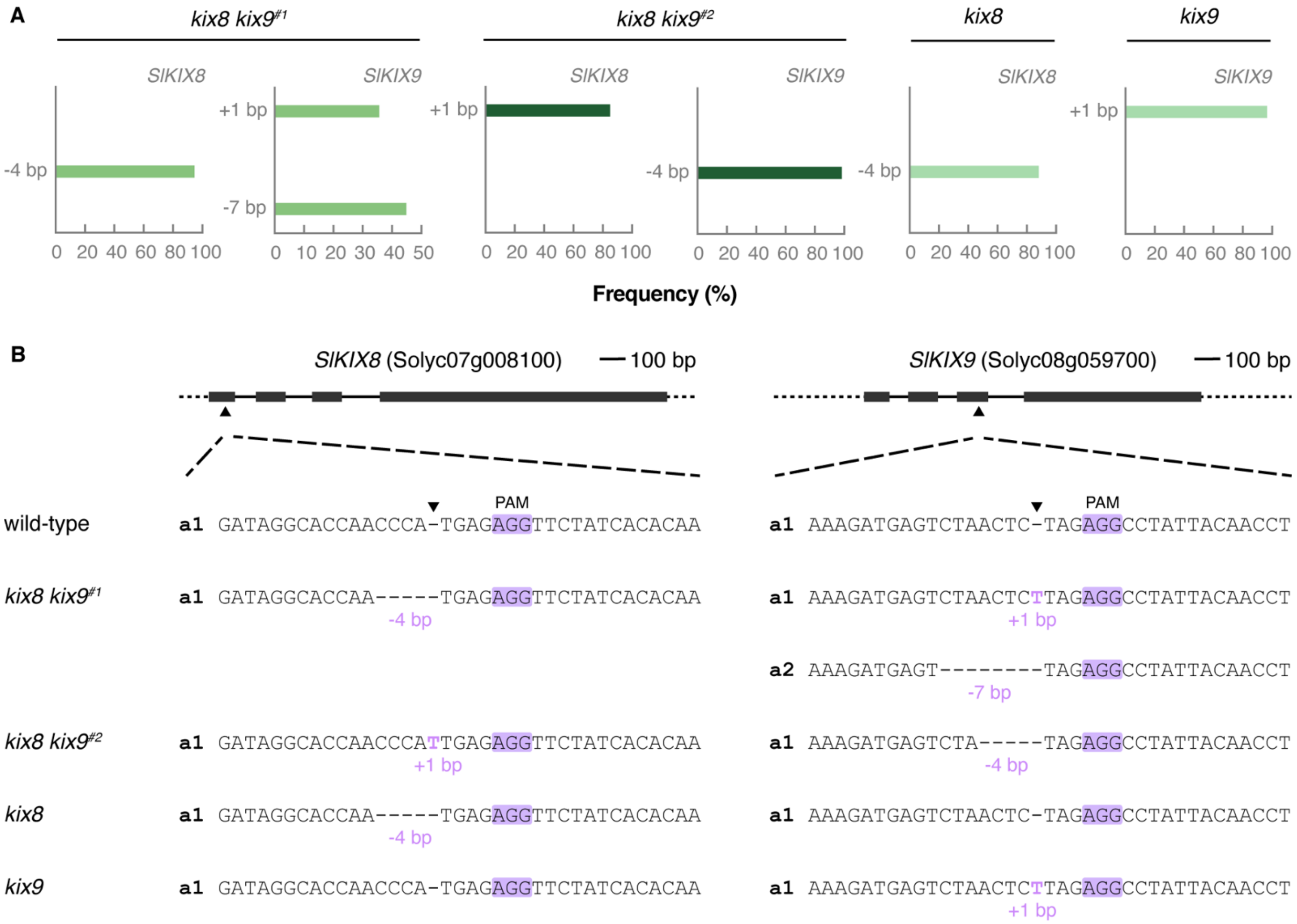


**Figure S4.** CRISPR-Cas9 mutations in double *kix8 kix9* (T1) and single *kix8*, and single *kix9* tomato knockout lines. (A) ICE analysis of genomic sites targeted by the guide RNAs. Targeted genomic regions were PCR amplified and sequenced by Sanger sequencing. Based on the sequence chromatograms, ICE analysis visualized the indel spectrum and calculated the frequency of each indel. (B) Schematic representation of *SlKIX8* and *SlKIX9* with location of the CRISPR-Cas9 cleavage sites. Dark grey boxes represent exons and solid lines represent introns. Cas9 cleavage sites for guide RNAs are indicated with arrowheads. Allele sequences are shown for two independent *kix8 kix9* lines, one *kix8* line, and one *kix9* line. Abbreviations: a, allele; PAM, protospacer adjacent motif.

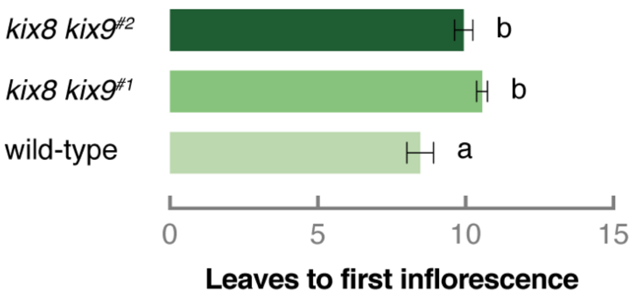


**Figure S5.** Tomato *kix8 kix9* plants display a delay in flowering time. Plants were grown in soil under 16:8 photoperiods with daytime and nighttime temperatures of 26–29°C and 18–20°C, respectively. Bars represent mean number of leaves produced before initiation of the first inflorescence. Error bars denote standard error (n = 15–16). Statistical significance was determined by ANOVA followed by Tukey post-hoc analysis ($P < 0.05$; indicated by different letters).

**Table S1.** Tomato *kix8 kix9* plants display a reduction in plant height.

| | wild-type | *kix8 kix9*$^{\#1}$ | *kix8 kix9*$^{\#2}$ |
|---|---|---|---|
| Height of the primary shoot (cm) | 12.33 ± 1.00[a] | 7.83 ± 0.47[b] | 7.83 ± 0.22[b] |
| Height of the main shoot (cm) | 20.33 ± 1.68[a] | 13.58 ± 0.84[b] | 12.67 ± 0.64[b] |
| Internode length (cm) | 2.50 ± 0.19[a] | 1.22 ± 0.09[b] | 1.19 ± 0.06[b] |

Plants were grown in soil for 4 months under 16:8 photoperiods with daytime and nighttime temperatures of 26–29°C and 18–20°C, respectively. Data are mean ± standard error ($n = 12$). Statistical significance was determined by ANOVA followed by Tukey post-hoc analysis ($P < 0.05$; indicated by different letters).

**Table S2.** Normalized expression of *SlKIX8*, *SlKIX9*, *SlPPD1*, *SlPPD2*, *SlDFL1*, *SlAHL17*, and *SlAP2d* in different tomato organs and developmental stages (cultivar Micro-Tom) used to generate heat maps in Figure 4, A and B.

| | | | Flower | | Fruit peel | | | | | Fruit flesh | | | | | Seed | | | | |
|---|---|---|---|---|---|---|---|---|---|---|---|---|---|---|---|---|---|---|---|
| | Root | Leaf | Bud | Petal | IG | MG | BR | OR | RR | IG | MG | BR | OR | RR | IG | MG | BR | OR | RR |
| *KIX8* | 0.040 | 0.022 | 0.061 | 0.079 | 0.026 | 0.011 | 0.013 | 0.006 | 0.0134 | 0.044 | 0.018 | 0.022 | 0.008 | 0.009 | 0.120 | 0.078 | 0.103 | 0.179 | 0.158 |
| *KIX9* | 0.053 | 0.008 | 0.021 | 0 | 0 | 0 | 0 | 0 | 0 | 0 | 0.001 | 0 | 0 | 0 | 0.001 | 0.001 | 0 | 0 | 0 |
| *PPD1* | 0.006 | 0.017 | 0.041 | 0.010 | 0.010 | 0.006 | 0.010 | 0 | 0 | 0.020 | 0.003 | 0.005 | 0 | 0.002 | 0.035 | 0.015 | 0.011 | 0.006 | 0.012 |
| *PPD2* | 0.098 | 0.035 | 0.090 | 0.039 | 0.020 | 0.034 | 0.060 | 0.050 | 0.080 | 0.040 | 0.020 | 0.037 | 0.029 | 0.061 | 0.066 | 0.069 | 0.074 | 0.053 | 0.082 |
| *DFL1* | 0.031 | 0.067 | 0.190 | 0.033 | 0.007 | 0.011 | 0 | 0 | 0 | 0.010 | 0.007 | 0.007 | 0.019 | 0.004 | 0.049 | 0.165 | 0.633 | 0.980 | 0.450 |
| *AHL17* | 0.123 | 0 | 0.098 | 0.043 | 0.010 | 0.065 | 0.040 | 0.110 | 0.310 | 0.010 | 0.017 | 0.015 | 0.041 | 0.253 | 0.071 | 0.158 | 0.533 | 0.925 | 0.405 |
| *AP2d* | 0.164 | 0.046 | 0.092 | 0.024 | 0.542 | 1.108 | 0.520 | 0 | 0 | 0.120 | 0.223 | 0.323 | 0.036 | 0.003 | 0.032 | 0.038 | 0.019 | 0.023 | 0.025 |

Expression data was obtained from TomExpress (Zouine *et al.*, 2017). Abbreviations: IG, immature green; MG, mature green; BR, breaker; OR, orange; RR, red ripe.

**Table S3.** Tomato *kix8 kix9* plants display a reduction in axillary shoot formation

| | wild-type | *kix8 kix9*[#1] | *kix8 kix9*[#2] |
|---|---|---|---|
| **Inflorescence parameters:** | | | |
| Main shoot | | | |
| Number of inflorescences | 2.75 ± 0.18[a] | 2.75 ± 0.18[a] | 2.83 ± 0.24[a] |
| Number of flowers per inflorescence | 14.24 ± 1.09[a] | 10.26 ± 0.93[b] | 7.36 ± 0.84[c] |
| Number of pollinated flowers per inflorescence | 8.16 ± 1.05[a] | 4.72 ± 0.88[b] | 3.14 ± 0.62[b] |
| Pollinated/total number of flowers per inflorescence (%) | 57.01 ± 7.04[a] | 42.25 ± 7.54[a] | 45.53 ± 6.32[a] |
| Axillary shoots | | | |
| Number of inflorescences | 5.17 ± 0.49[a] | 2.25 ± 0.37[b] | 2.75 ± 0.13[b] |
| **Fruit parameters:** | | | |
| Main shoot and axillary shoots | | | |
| Green fruit biomass (g) | 0.93 ± 0.60[a] | 1.38 ± 1.11[a] | 1.59 ± 0.99[a] |
| Breaker–orange fruit biomass (g) | 1.50 ± 0.77[a] | 2.11 ± 0.89[a] | 3.90 ± 1.21[b] |
| Red fruit biomass (g) | 2.01 ± 0.96[a] | 3.68 ± 1.16[b] | 4.65 ± 1.27[c] |
| Main shoot | | | |
| Breaker–red fruit pericarp thickness (mm) | 1.55 ± 0.09[a] | 2.86 ± 0.12[b] | 3.23 ± 0.09[c] |
| Breaker–red fruit pericarp thickness/radius (%) | 19.46 ± 0.42[a] | 29.18 ± 0.78[b] | 29.30 ± 0.37[b] |
| **Fruit yield parameters:** | | | |
| Main shoot | | | |
| Number of green fruits | 1.58 ± 0.63[a] | 4.58 ± 0.74[b] | 1.50 ± 0.38[a] |
| Number of breaker–orange fruits | 1.75 ± 0.80[a] | 1.58 ± 0.45[a] | 0.75 ± 0.30[a] |
| Number of red fruits | 18.92 ± 1.96[a] | 8.92 ± 0.53[b] | 8.00 ± 0.72[b] |
| Total fruit number | 22.25 ± 2.06[a] | 15.08 ± 1.23[b] | 10.25 ± 0.76[b] |
| Green fruit yield (g) | 2.09 ± 1.06[a] | 6.81 ± 1.70[b] | 2.50 ± 0.79[ab] |
| Breaker–orange fruit yield (g) | 1.42 ± 0.63[a] | 3.51 ± 1.02[a] | 2.76 ± 1.11[a] |
| Red fruit yield (g) | 39.25 ± 4.71[a] | 31.75 ± 2.27[a] | 35.50 ± 1.91[a] |
| Total fruit yield (g) | 42.77 ± 4.99[a] | 42.07 ± 3.06[a] | 40.76 ± 2.05[a] |
| Axillary shoots | | | |
| Number of green fruits | 16.00 ± 3.57[a] | 0.42 ± 0.19[b] | 0.25 ± 0.13[b] |
| Number of breaker–orange fruits | 5.42 ± 1.37[a] | 0.50 ± 0.50[b] | 0.00 ± 0.00[b] |
| Number of red fruits | 1.67 ± 0.57[a] | 0.42 ± 0.34[a] | 0.92 ± 0.26[a] |
| Total fruit number | 23.08 ± 4.56[a] | 1.33 ± 0.91[b] | 1.17 ± 0.27[b] |
| Green fruit yield (g) | 12.25 ± 1.33[a] | 0.51 ± 0.27[b] | 0.28 ± 0.19[b] |
| Breaker–orange fruit yield (g) | 8.00 ± 2.00[a] | 1.25 ± 1.25[b] | 0.00 ± 0.00[b] |
| Red fruit yield (g) | 3.58 ± 1.36[a] | 0.92 ± 0.75[a] | 4.00 ± 1.27[a] |
| Total fruit yield (g) | 23.83 ± 2.78[a] | 2.68 ± 2.17[b] | 4.28 ± 1.29[b] |
| Main shoot and axillary shoots | | | |
| Number of green fruits | 17.58 ± 3.54[a] | 5.00 ± 0.72[b] | 1.75 ± 0.39[b] |
| Number of breaker–orange fruits | 7.17 ± 1.78[a] | 2.08 ± 0.56[b] | 0.75 ± 0.30[b] |
| Number of red fruits | 20.58 ± 1.61[a] | 9.33 ± 0.43[b] | 8.92 ± 0.69[b] |
| Total fruit number | 45.33 ± 3.75[a] | 16.42 ± 0.96[b] | 11.42 ± 0.71[b] |
| Green fruit yield (g) | 14.34 ± 1.52[a] | 7.32 ± 1.75[b] | 2.78 ± 0.93[b] |
| Breaker–orange fruit yield (g) | 9.42 ± 2.21[a] | 4.76 ± 1.34[ab] | 2.76 ± 1.11[b] |
| Red fruit yield (g) | 42.83 ± 3.72[a] | 32.67 ± 1.87[b] | 39.50 ± 2.10[ab] |
| Total fruit yield (g) | 66.60 ± 3.18[a] | 44.75 ± 2.37[b] | 45.04 ± 2.24[b] |
| **Seed yield parameters:** | | | |
| Main shoot | | | |
| Number of seeds in red fruit | 11.42 ± 8.54[a] | 6.92 ± 5.30[b] | 5.95 ± 4.73[b] |
| Seed area in red fruit ($mm^2$) | 5.98 ± 0.84[ab] | 6.11 ± 1.45[a] | 5.50 ± 1.35[b] |

Plants were grown in soil for 3.5–4.5 months under 16:8 photoperiods with daytime and nighttime temperatures of 26–29°C and 18–20°C, respectively. Inflorescence parameters were measured and fruits were harvested from each genotype when the ratio of ripe to unripe fruits on the main shoot was 60–85%. Data are mean ± standard error (n = 12). Statistical significance was determined by ANOVA followed by Tukey post-hoc analysis ($P < 0.05$; indicated by different letters).

**Table S4.** Oligonucleotides used in this study.

| Oligonucleotide | Sequence (5'–3') | Orientation | Description | SolycID |
|---|---|---|---|---|
| Oligonucleotides for Y2H/Y3H constructs: | | | | |
| LAPAU2860 | GGGGACAAGTTTGTACAAAAAAGCAGGCTCCATGCCTAGACCAGGACCCAG | Forward | amplification of *KIX8* | Solyc07g008100.2 |
| LAPAU2994 | GGGGACCACTTTGTACAAGAAAGCTGGGTCTCMCAAACCTGGCCTTTTCATTTG | Reverse | | |
| LAPAU2862 | GGGGACAAGTTTGTACAAAAAAGCAGGCTCCATGCCTAAATCTACAAGAGC | Forward | amplification of *KIX9* | Solyc08g059700.1 |
| LAPAU2863 | GGGGACCACTTTGTACAAGAAAGCTGGGTCTCMGGACTTGAATTTGTTAAAATG | Reverse | | |
| LAPAU2856 | GGGGACAAGTTTGTACAAAAAAGCAGGCTCCATGCCGCCGGAAGAAACAG | Forward | amplification of *PPD1* | Solyc06g084120.2 |
| LAPAU2857 | GGGGACCACTTTGTACAAGAAAGCTGGGTCTCMCTTTCTAACATCTCTGTC | Reverse | | |
| LAPAU2858 | GGGGACAAGTTTGTACAAAAAAGCAGGCTCCATGTCGCTGGAACAAACTG | Forward | amplification of *PPD2* | Solyc09g065630.2 |
| LAPAU2859 | GGGGACCACTTTGTACAAGAAAGCTGGGTCTCMCTCTTTACCATCTTTG | Reverse | | |
| COMBI6198 | GGGGACAAGTTTGTACAAAAAAGCAGGCTCCATGTCGTCTTCACAATCACCACCATC | Forward | amplification of *SAP* | Solyc05g041220.2 |
| COMBI6199 | GGGGACCACTTTGTACAAGAAAGCTGGGTTCACTATTGTGCACCAAAGTCCCACAAATG | Reverse | | |
| Oligonucleotides for CRISPR-Cas9 constructs: | | | | |
| LAPAU2582 | ATTGATAGGCACCAACCCATGAG | Forward | *KIX8* gRNA target site | Solyc07g008100.2 |
| LAPAU2583 | AAACCTCATGGGTTGGTGCCTAT | Reverse | | |
| LAPAU2580 | ATTGAAAGATGAGTCTAACTCTAG | Forward | *KIX9* gRNA target site | Solyc08g059700.1 |
| LAPAU2581 | AAACCTAGAGTTAGACTCATCTTT | Reverse | | |
| Oligonucleotides for the identification of CRISPR-Cas9 mutants: | | | | |
| LAPAU3075 | TCCCTCATCAGATCCACCTC | Forward | amplification of *Cas9* | – |
| LAPAU3076 | CTGAAACCTGAGCCTTCTGG | Reverse | | |
| LAPAU2783 | CCCCTCCAAAACACTCATGT | Forward | amplification of *KIX8* gRNA target region | Solyc07g008100.2 |
| LAPAU2784 | GAGCAGTACAAATGAGCAGCA | Reverse | | |
| LAPAU2785 | GCTGAAGAAATTATGTATTCCAAAGC | Forward | amplification of *KIX9* gRNA target region | Solyc08g059700.1 |
| LAPAU2786 | CCCGAGAAGTTTCACTCGAA | Reverse | | |
| Oligonucleotides for gene expression analysis by qPCR: | | | | |
| COMBI5428 | CCTCCGTTGTGATGTAACTGG | Forward | amplification of *CAC* | Solyc08g006960.2 |
| COMBI5429 | ATTGGTGGAAAGTAACATCATCG | Reverse | | |
| COMBI5416 | ATGGAGTTTTTGAGTCTTCTGC | Forward | amplification of *TIP41* | Solyc10g049850.1 |
| COMBI5417 | GCTGCGTTTCTGGCTTAGG | Reverse | | |
| COMBI7162 | ACCATCGAAGAGTCTCTCAACAGC | Forward | amplification of *DFL1* | Solyc07g063850.2 |
| COMBI7163 | CAATGGATTTGTCTGAGGCACGAC | Reverse | | |
| COMBI7168 | CTGTCATTTGCCGTCGGATGTG | Forward | amplification of *AHL17* | Solyc04g076220.2 |
| COMBI7169 | AGTAAGGCGGTGGTTGTGGTTG | Reverse | | |
| COMBI7158 | TGCATAGTCAGGTCGGAACAACG | Forward | amplification of *AP2d* | Solyc11g072600.1 |
| COMBI7159 | TGGTAGCCGGAGTTGAGAATCC | Reverse | | |
| COMBI7188 | AGGCTGTGTCTACCAGCAAAGAC | Forward | amplification of *KIX8* | Solyc07g008100.2 |
| COMBI7189 | TTGCAACCCGGAGTGACTGTTG | Reverse | | |
| COMBI7190 | AGACACCAACCAATCAGAGGTTCC | Forward | amplification of *KIX9* | Solyc08g059700.1 |
| COMBI7191 | TGCTGAGCCATGAACCTCATTCAC | Reverse | | |
